# *Myrmecocystus* honeypot ants have a species-specific resident gut microbiome

**DOI:** 10.64898/2026.04.07.717087

**Authors:** DiemQuynh V. Nguyen, Charlotte B. Francoeur, Bianca R. Nogueira, Indira Sawh, Michele Lanan, Lily Khadempour

## Abstract

*Myrmecocystus* honeypot ants rely on specialized workers, known as repletes, to store dissolved carbohydrates in their crops. The repletes store this liquid for many months at a time without spoilage. When resources are scarce, repletes redistribute the stored nutrients to their colony members via trophallaxis. While we suspect that the gut microbiome of honeypot ants may aid in spoilage prevention, we must first characterize it. Here, we used 16S rRNA gene sequencing to determine the microbial community composition across six *Myrmecocystus* honeypot ant species, sampling multiple colonies, castes, and organs across four years. We found microbiome community composition was strongly shaped by species, with variation between colonies in *M. arenarius*, *M. depilis*, and *M. mexicanus.* Differences between forager and replete castes were observed in *M. arenarius*, *M. flaviceps,* and *M. mexicanus.* Organ-level differences were observed in the crop and midgut in *M. mexicanus*. Replete crops of *M. mexicanus* and *M. depilis* were enriched in *Fructilactobacillus,* other lactic acid bacteria, and acetic acid bacteria. These bacteria turn sugars into organic acids that lower pH and can inhibit other microbes. Halophiles were prominent in the guts of *M. flaviceps* and *M. wheeleri*, which may reflect the high osmotic stress of concentrated sugars in stored fluids. In this study, we demonstrate that *Myrmecocystus* ants host species-specific gut microbiomes. While much work remains to understand the relationship between the host ant and its gut microbiome, the repeated presence of acid-producing and stress-tolerant bacteria suggests honeypot ants’ gut microbiome could potentially keep crop contents stable.

**Importance:** Previous work showed that fluid stored by Australian honeypot ants has antimicrobial activity, likely influenced by the ants’ gut microbiome. Our study is the first to characterize the gut microbiome of North American honeypot ant species in the genus *Myrmecocystus.* Our work shows that microbiome composition varies substantially between species, with distinct enrichments of lactic acid bacteria, acetic acid bacteria, or halophiles. These findings suggest that host ecology and environmental conditions, such as diet and the crop’s unique storage environment, can help structure microbial communities. This system provides a framework to examine how host-associated environments shape microbiome composition and function. Understanding microbiome variation in honeypot ants creates opportunities for future work to investigate how microbial communities contribute to the long-term preservation of nutrient-rich resources, analogous to human food preservation and fermentation processes. Our work establishes *Myrmecocystus* as a model for studying host-microbe symbiosis in extreme nutritional environments, enabling future hypothesis-driven research into microbial function, community assembly, and the evolutionary dynamics of host-associated microbiomes.

## Introduction

Many animals, particularly those with specialized diets, rely on their microbiomes to supplement essential nutrients and provide beneficial services to the host (1). Compared to vertebrates, insects provide more tractable systems for studying host-microbiome relationships because they tend to have simpler gut morphology and gut microbiome communities. Host-associated microbiomes can contribute to a range of tasks, including nitrogen fixation and recycling (e.g., *Cephalotes* turtle ants), amino acid production (e.g., *Onthophagus taurus* dung beetles), toxin remediation (e.g., *Apis mellifera* honeybees), and the breakdown of complex plant polysaccharides the host cannot digest independently (e.g., *Nasuititermes* termites) (2–7). However, not all insects rely on a gut microbiome. For example, despite their herbivorous diet, *Lepidoptera* caterpillars, which have a transient microbiome, can meet their nutritional demands through host-encoded digestive enzymes and intrinsic mechanisms to degrade or tolerate plant toxins (8).

Honeypot ants represent a compelling system for investigating animal-associated gut microbiomes. This convergently evolved group comprises at least eight independently evolved genera spanning three Formicidae subfamilies (9, 10). In this study, we focus on the North American honeypot ants in the genus *Myrmecocystus*. This genus began diversifying approximately 14 million years ago, coinciding with the aridification of the southwestern United States and northern Mexico (11). The formation of new desert habitats created ecological niches that possibly shaped the evolution of this group (11). To overcome irregular resource availability, honeypot ants have evolved specialized workers, known as repletes, that act as living food storage units to sustain the colony during dry, nutrient-limited seasons (10).

Honeypot ant foragers collect extrafloral nectar, honeydew, and insect fluids to provide them to the repletes through trophallaxis, a regurgitative food-sharing process common to social insects (12). As repletes accumulate the collected resources, the crops become enlarged, and the intersegmental membranes of the gaster undergo extensive stretching (Fig. 1B, D-E) (13). While the basic structure of adult ant digestive tracts is conserved across all ants, honeypot ants can distend their crop far beyond the limits observed in non-replete ants, facilitating long-term storage of liquid resources (14). A study by Dong et al. (2023) found the liquid stored in Australian honeypot ants, *Camponotus inflatus*, has antimicrobial properties (15). Based on whole-ant sequencing, they concluded that the microbiome was dominated by the endosymbiotic bacterial genus *Candidatus Blochmannia* and the fungal genus *Neocelosporium* (15).

**Figure 1.**
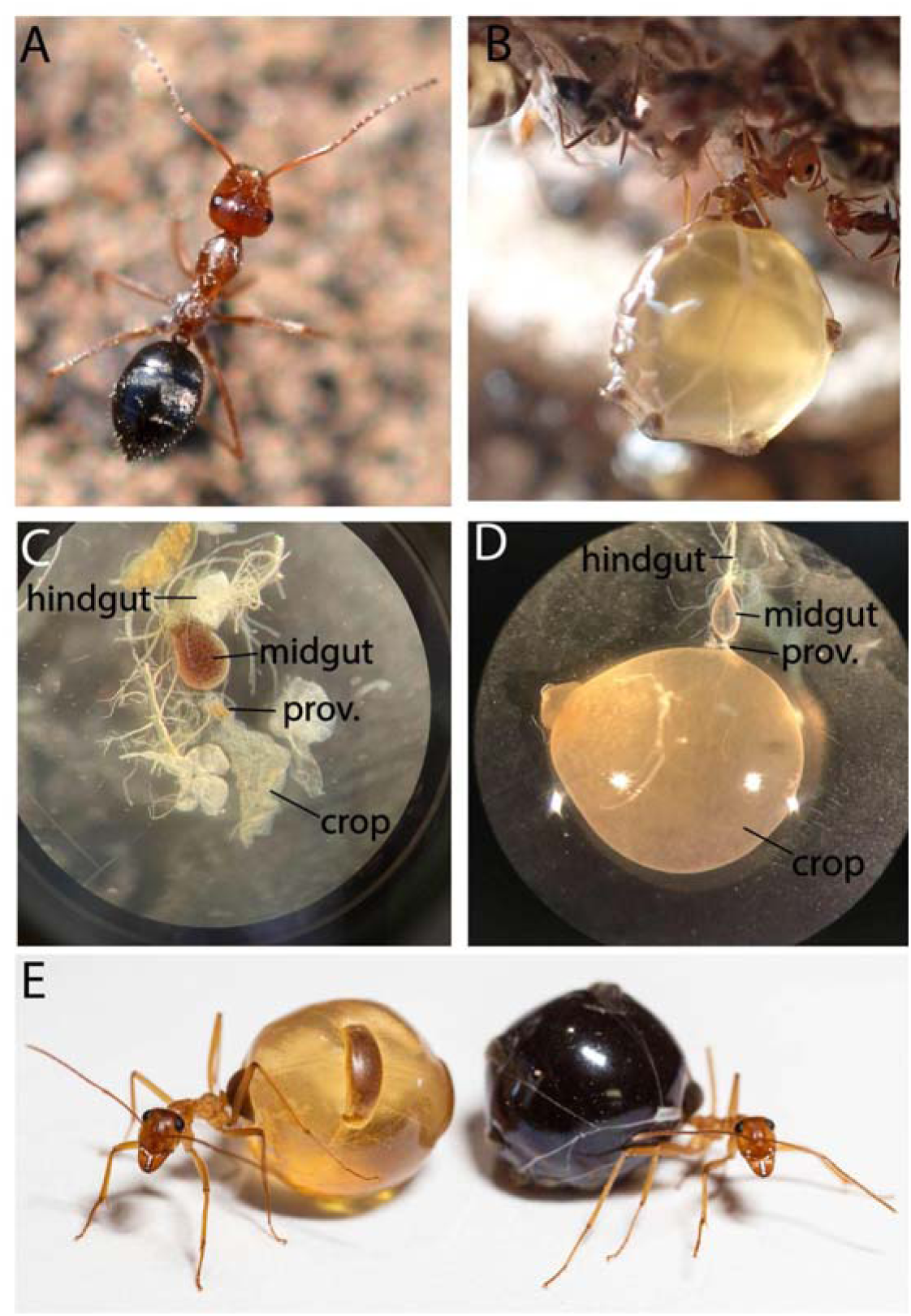
Honeypot ant colonies contain two castes, workers (A) and repletes (B). Compared to the crops of workers (C), the crops of repletes can expand to huge proportions as they are used to store fluids. (E) Depending on the resources collected, repletes within the same colony or chamber can have abdomens of different colors. (Photos credits: A: D. Caroll from buigguide.net, B: M.P. Meurville, C: I. Sawh, D: C. Francoeur E: M. Lanan).

*Blochmannia* is a well-known endosymbiont of *Camponotus* ants, residing within bacteriocytes associated with the midgut, the organ where most digestion occurs (16). Since *Blochmannia* is restricted to midgut tissues, we do not think it contributes to antimicrobial activity. Instead, we would expect that if a microbe were responsible for antimicrobial properties, it would reside within the crop rather than in bacteriocytes associated with the midgut (16, 17). In our work, instead of characterizing the microbiome of the whole gaster or the whole ant, we dissect the ants and characterize the contents of the crop and midgut separately. We hypothesize that the gut microbiome of *Myrmecocystus* honeypot ants contributes to antimicrobial activity within their sugar-rich crop, thereby limiting the growth of pathogenic bacteria and fungi and preventing spoilage. However, before addressing potential functions, it is necessary to first characterize the gut microbiome of *Myrmecocystus* species. We intend to determine if *Myrmecocystus* has a resident gut microbiome (i.e., if it has a microbiome that is consistent across individuals within a colony and within a species).

In this study, we characterize the gut microbiome of six *Myrmecocystus* honeypot ant species: *M. arenarius, M. depilis, M. flaviceps, M. mexicanus, M. mimicus,* and *M. wheeleri*. We use community amplicon sequencing to determine whether a consistent resident gut microbiome exists. We compare the gut microbiome across scales: between different species within the genus, between colonies of the same species, between castes within a colony, and between digestive organs within individual ants. If these ants possess a resident gut microbiome, we expect to observe the same community members in the same organs within a species, and potentially across the genus, over multiple colonies and years. Establishing whether *Myrmecocystus* hosts a resident gut microbiome is an important first step in understanding the relationship between these ants and their associated microbes.

## Results

### Species Differences

We characterized the gut microbiomes of a total of 321 samples across six species: *M. arenarius* (n = 35), *M. depilis* (n = 39), *M. flaviceps* (n = 27), *M. mexicanus* (n = 166), *M. mimicus* (n = 26), and *M. wheeleri* (n = 28). We found microbiome composition differed significantly among the six *Myrmecocystus* species in both organs (permutational multivariate analysis of variance (PERMANOVA) species crop: R^2^ = 0.699, p ≤ 0.001; species midgut: R^2^ = 0.577, p ≤ 0.001; Table S4; Fig. 2; Fig. 3). However, we found significant differences in multivariate dispersion among species in both organs (PERMDISP crop F = 17.93, p ≤ 0.001; midgut: F = 4.92, p ≤ 0.002; Table S4), suggesting that dispersion may contribute to the observed species-level variation in microbiome composition.

**Figure 2.**
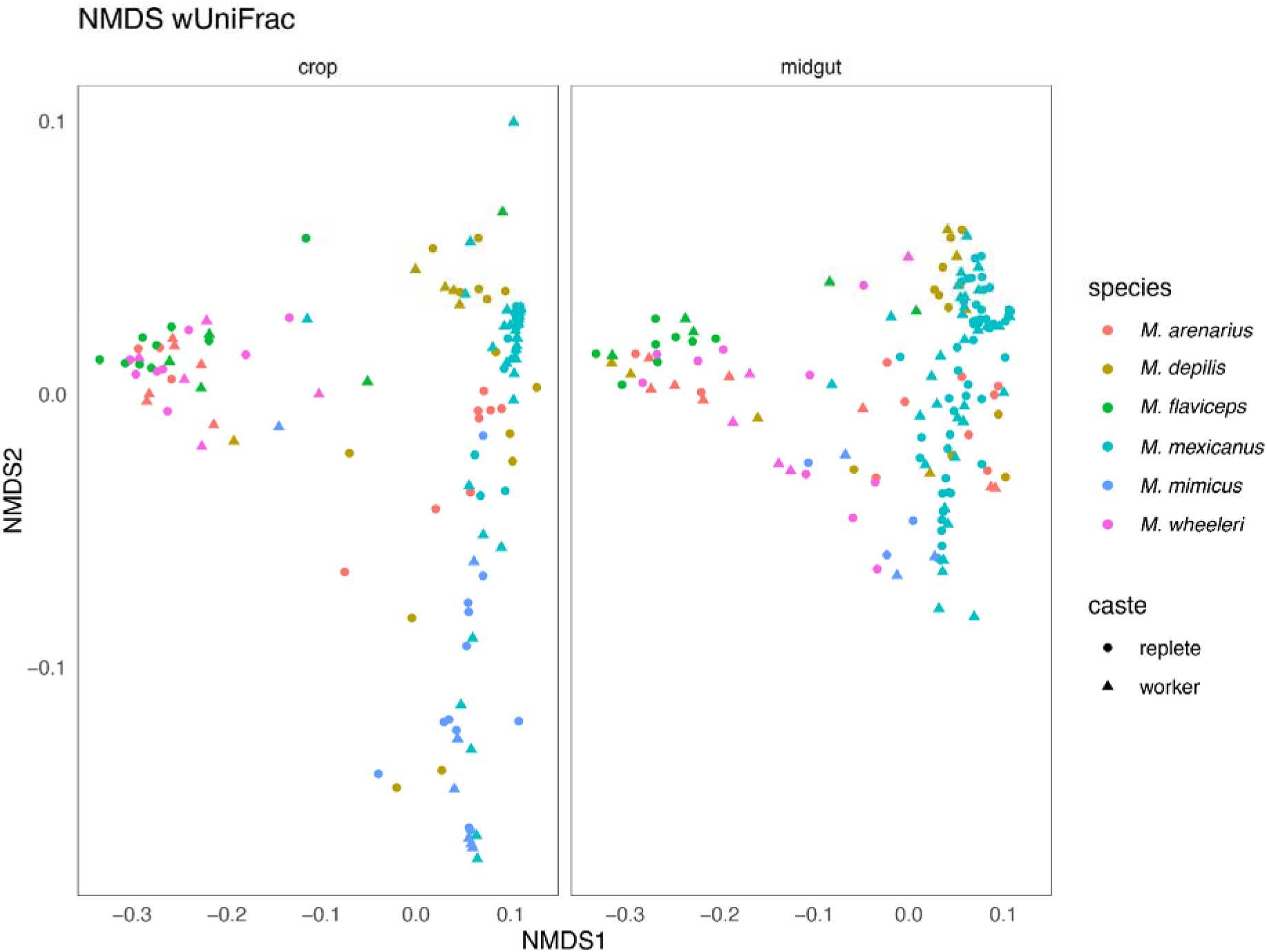
Non-metric Multidimensional Scaling (NMDS) ordination based on weighted UniFrac distances visualizes differences in microbial community composition across *Myrmecocystus* species, caste, and organ. Each point represents individual samples, species indicated by color, caste by shape, with panels corresponding to crop or midgut samples. The ordination includes 321 organ samples across the six species: *M. arenarius* (n = 35), *M. depilis* (n = 39), *M. flaviceps* (n = 27), *M. mexicanus* (n = 166), *M. mimicus* (n = 26), and *M. wheeleri* (n = 28). Microbial communities show species-associated clustering. However, the amount of variation within each species also differed significantly (PERMDISP p = 0.001), indicating that some species had more variable microbiome communities than others. Crop and midgut communities also differed significantly overall after accounting species identity, although organ explained a small proportion of variation than species (PERMANOVA organ R^2^ = 0.030, p = 0.000999).

**Figure 3.**
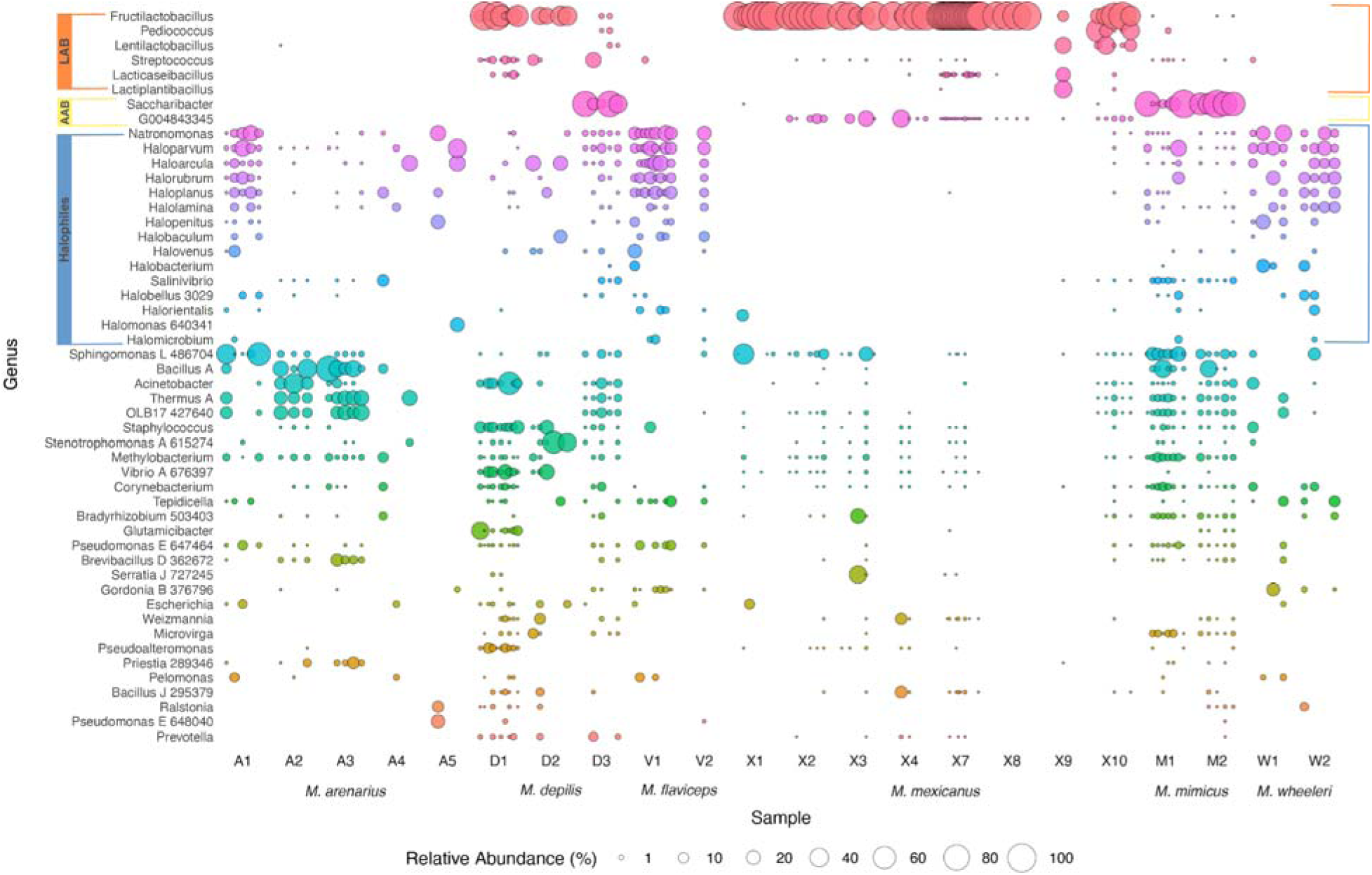
The bubble plot depicts the relative abundance (as a percentage) of bacterial genera in replete crop samples across *Myrmecocystus* species. Bubble size indicates relative abundance, and color signify genus. The total sample size was 105 crop samples: *M. arenarius* (n = 11), *M. depilis* (n = 16), *M. flaviceps* (n =9), *M. mexicanus* (n = 49), *M. mimicus* (n=12), and *M. wheeleri* (n = 8). Genus names are grouped by lactic acid bacteria (LAB = orange), acetic acid bacteria (AAB = yellow), and halophilic archaea and bacteria (blue). Letter prefixes on the x axis denote species and their colony, different numbers within the same prefix represent distinct colonies of the same species (e.g., A1 and A2 are separate *M. arenarius* colonies). The microbial communities in replete crops exhibit species-specific structuring. In particular *M. mexicanus* replete crops are strongly dominated by *Fructilactobacillus*, whereas other species show more variable community composition.

We found significant differences in Shannon diversity across the six species (Kruskal-Wallis chi-squared = 58.922, df = 5, p = 2.03e^-11^), even after accounting for caste, colony, and individual ant identity as random effects (GLMM, Wald χ² = 29.89, df = 5, p = 1.55 × 10 ; Table S9). *Myrmecocystus mexicanus* had significantly lower Shannon diversity than *M. arenarius, M. depilis, M. flaviceps*, and *M. wheeleri,* but did not differ significantly from *M. mimicus* (Table S2). Species differences in Shannon diversity also varied between the crop and midgut (GLMM, Wald χ² = 101.06, df = 5, p < 0.001), where *M. mexicanus* had significantly lower crop diversity compared to all species (p < 0.0001; Table S11). No significant pairwise differences were detected in midgut Shannon diversity across species (Turkey-adjusted p > 0.05; Table S11). Among the 105 replete crop samples examined across the six species, *M. mexicanus* replete crops (n = 49) were dominated by *Fructilactobacillus*, with the three most abundant ASVs belonging to this genus (ASV1, ASV5, and ASV12) (Fig. 3, Fig. S2). In contrast, *Fructilactobacillus* was largely absent from *M. arenarius* and *M. wheeleri* replete gut microbiomes (Fig. 3, Fig. S2). Instead, these two species showed a greater abundance of halophilic taxa, such as *Haloparvum* (Fig. 3, Fig. S2).

Overall, our results suggest that species identity plays a substantial role in microbiome variation, particularly in the crop. Notably, *M. mexicanus* had a less diverse crop microbiome dominated by *Fructilactobacillus*, whereas other species generally exhibited more diverse crop microbiomes (Fig. S2).

### Colony Difference Within Species

When we examined whether microbiome composition differed among colonies within each *Myrmecocystus* species, we found colony-associated differences in some, but not all, species (Table S4). *Myrmecocystus arenarius* (4 colonies; crop n = 11, midgut n = 10)*, M. depilis* (3 colonies; crop n = 16, midgut n = 10), and *M. mexicanus* (8 colonies; crop n = 49, midgut n = 51) showed significant differences among colonies in both crop and midgut communities (UniFrac distance, PERMANOVA: p ≤ 0.001; R^2^ = 0.229 to 0.702; Table S4). ANOSIM also identified significant colony-associated differences in microbiome composition in these three species (Table S4). However, we found no significant colony differences in *M. flaviceps* (2 colonies; crop n = 9, midgut n = 8)*, M. mimicus* (2 colonies; crop n = 12, midgut n = 4), and *M. wheeleri* (2 colonies; crop n = 8, midgut n = 10) (PERMANOVA and ANOSIM p > 0.05; Table S4).

We also examined whether microbiome composition varied across collection times in *M. mexicanus* and *M. depilis* (Table S5). In *M. mexicanus,* collection time was not significantly associated with differences in replete crop microbiome composition (PERMANOVA R^2^ = 0.083, p = 0.0749), whereas replete midgut microbiome differed significantly among collection times (PERMANOVA R^2^ = 0.537, p ≤ 0.001). Multivariate dispersion did not differ significantly among collection times in either organ (PERMDISP crop p = 0.121; midgut p = 0.749). Pairwise comparisons further showed significant differences in midgut microbiome composition among all three collection times, whereas crop differences were more limited.

Despite variation across collection times, *Fructilactobacillus* remained a dominant genus in *M. mexicanus* crop, with ASV1 present in all samples (100% prevalence). However, the relative abundance of other bacterial taxa varied across collection times, including *Acetobacteraceae* G004843345 (ASV6), *Weizmannia* (ASV21), and lactic acid bacteria such as *Pediococcus* (ASV23) and *Lentilactobacillus* (ASV42).

In *M. depilis,* replete crop communities differed significantly between 2023 and 2025 collections (PERMANOVA R^2^ = 0.471, p ≤ 0.001; Table S5). In contrast, we found no significant differences in midgut microbiome composition between collection years (p = 0.0979), although this comparison was limited by having only one midgut sample from July 2025. In 2023, dominant crop ASVs included *Fructilactobacillus* (ASV1), *Acinetobacter* (ASV73), and *Vibrio* A 676397 (ASV7), whereas the 2025 samples showed greater relative abundance of *Saccharibacter* (ASV11) and other taxa.

Comparisons of GLMMs with and without colony identity as a random effect showed that including colony significantly improved model fit (likelihood-ratio χ² = 32.68, df = 1, p < 0.001). The colony-level intraclass correlation coefficient (ICC) was 0.150 compared with 0.033 for individual ant identity. Fixed effects explained 40.9% of the variance in Shannon diversity (marginal R^2^ = 0.409), while the full model including random effects explained 51.7% (conditional R^2^ = 0.517). These findings suggest that colony identity contributes to variation in Shannon diversity beyond the effects of species, organ, and caste.

Overall, our results suggest that colony-associated differences in microbiome composition occur in some, but not all, *Myrmecocystus* species. We also found differences across collection times in some species and organs, suggesting that temporal variation may contribute to the observed microbiome patterns. However, these patterns should also be considered in the context of uneven sampling among colonies and collection times.

### Caste Differences (Replete vs. Worker)

Microbiome composition did not differ significantly between repletes (n = 198 organ samples) and workers (n = 123 organ samples) overall (PERMANOVA, crop p = 0.142; midgut p = 0.274; Table S4). However, when we examined caste differences in each species separately, crop microbiome composition differed significantly between castes in *M. flaviceps* (R^2^ = 0.236, p = 0.042), *M. mexicanus* (R^2^ = 0.140, p ≤ 0.001), and *M. arenarius* (R^2^ = 0.368, p = 0.009). These caste-associated differences in crop microbiome composition were also reflected in the relative abundance of dominant bacterial taxa. For example, in *M. mexicanus, Fructilactobacillus* dominated the crop microbiome of both castes, but worker crops also showed differences in the relative abundance of *Acetobacteraceae* G004843345 (ASV6 and ASV19). In contrast, we found no significant caste differences in crop microbiome composition in *M. depilis*, *M. wheeleri*, and *M. mimicus*. Multivariate dispersion did not differ significantly between replete and worker crop communities in *M. flaviceps* or *M. arenarius* crop communities (Table S4). However, we found significant differences in multivariate dispersion between *M. mexicanus* replete and worker crop communities (PERMDISP F = 25.51, p = 0.001; Table S4), suggesting that differences in within-caste variability may contribute to the observed PERMANOVA results.

Shannon diversity did not differ significantly between repletes (n = 198) and workers (n = 123) after accounting for species, organ, and individual ant identity (GLMM, Wald χ² = 2.80, df = 1, p = 0.0943). Although the estimated effect indicated slightly higher Shannon diversity in workers than repletes (β = 0.135), this difference was not statistically significant. However, when we examined caste differences within each species using pairwise Wilcoxon rank-sum tests, we found workers had significantly higher Shannon diversity than repletes in *M. arenarius* (FDR-adjusted p = 0.0264), *M. flaviceps* (FDR-adjusted p = 0.00453), and *M. mexicanus* (FDR-adjusted p = 0.00453). In contrast, we found no significant caste differences in Shannon diversity in *M. depilis*, *M. mimicus*, or *M. wheeleri* (FDR-adjusted p > 0.05) (Table S6).

Overall, our results suggest that caste plays a limited role in structuring gut microbiomes across *Myrmecocystus* species. Although crop microbiome composition differed between repletes and workers in *M. arenarius, M. flaviceps,* and *M. mexicanus*, we found no significant differences in midgut microbiome composition between castes across the six species.

### Organ Differences (Crop vs. Midgut)

We found microbiome composition differed significantly between the crop (n = 167) and midgut (n = 154), after accounting for species identity (PERMANOVA R^2^ = 0.030, p ≤ 0.001; Table S4). Multivariate dispersion did not differ significantly between organs overall (PERMDISP F = 0.40, p = 0.538), suggesting that differences in dispersion did not drive the overall organ-associated differences in microbiome composition.

When we examined organ differences in each species separately, we found significant differences in microbiome composition between the crop and midgut in *M. mexicanus* (R^2^ = 0.223, p ≤ 0.001), *M. mimicus* (R^2^ = 0.230, p = 0.002997), and *M. wheeleri* (R^2^ = 0.233, p = 0.001998). However, no significant organ effect was detected in *M. depilis, M. flaviceps*, or *M. arenarius* (Table S4). We also found significant differences in multivariate dispersion between the crop and midgut in *M. mexicanus* (PERMDISP F = 28.09, p = 0.001), suggesting that dispersion may contribute to the observed compositional differences between organs. However, multivariate dispersion did not differ significantly between organs in *M. mimicus* or *M. wheeleri*.

Shannon diversity differed significantly between the crop (n = 105) and midgut (n = 93) of repletes across the six species (GLMM, Wald χ² = 113.44, df = 1, p < 0.001). Within-species comparisons indicated significantly higher Shannon diversity in the midgut (2.572 ± 0.140 SE) than in the crop (0.943 ± 0.139 SE; p < 0.0001) for *M. mexicanus.* A significant difference was also detected in *M. mimicus*, with higher Shannon diversity in the midgut (2.558 ± 0.271 SE) than in the crop (1.965 ± 0.273 SE; p = 0.0058). In contrast, we did not find any significant differences in Shannon diversity between organs in *M. arenarius*, *M. depilis*, *M. flaviceps*, or *M. wheeleri* (all p > 0.05; Table S10). Consistent with the GLMM results, Wilcoxon rank-sum tests also detected higher Shannon diversity in *M. mexicanus* replete midgut samples (n = 51) than in the crop (n = 49) (FDR-adjusted p = 9.55e^-13^; Fig. 4). Similar patterns were seen for Inverse Simpson diversity and observed richness, with *M. mexicanus* midgut samples showing significantly higher alpha diversity than crop samples (FDR-adjusted p < 0.001; Fig. S3). In contrast, we found no significant differences in diversity between organs in other species (all FDR-adjusted p > 0.05) (Fig. 4; Fig. S3). Together, these findings indicate that organ-associated differences in Shannon diversity were particularly pronounced in *M. mexicanus*. GLMM also identified a significant difference in *M. mimicus* that was not detected by the Wilcoxon analysis.

**Figure 4.**
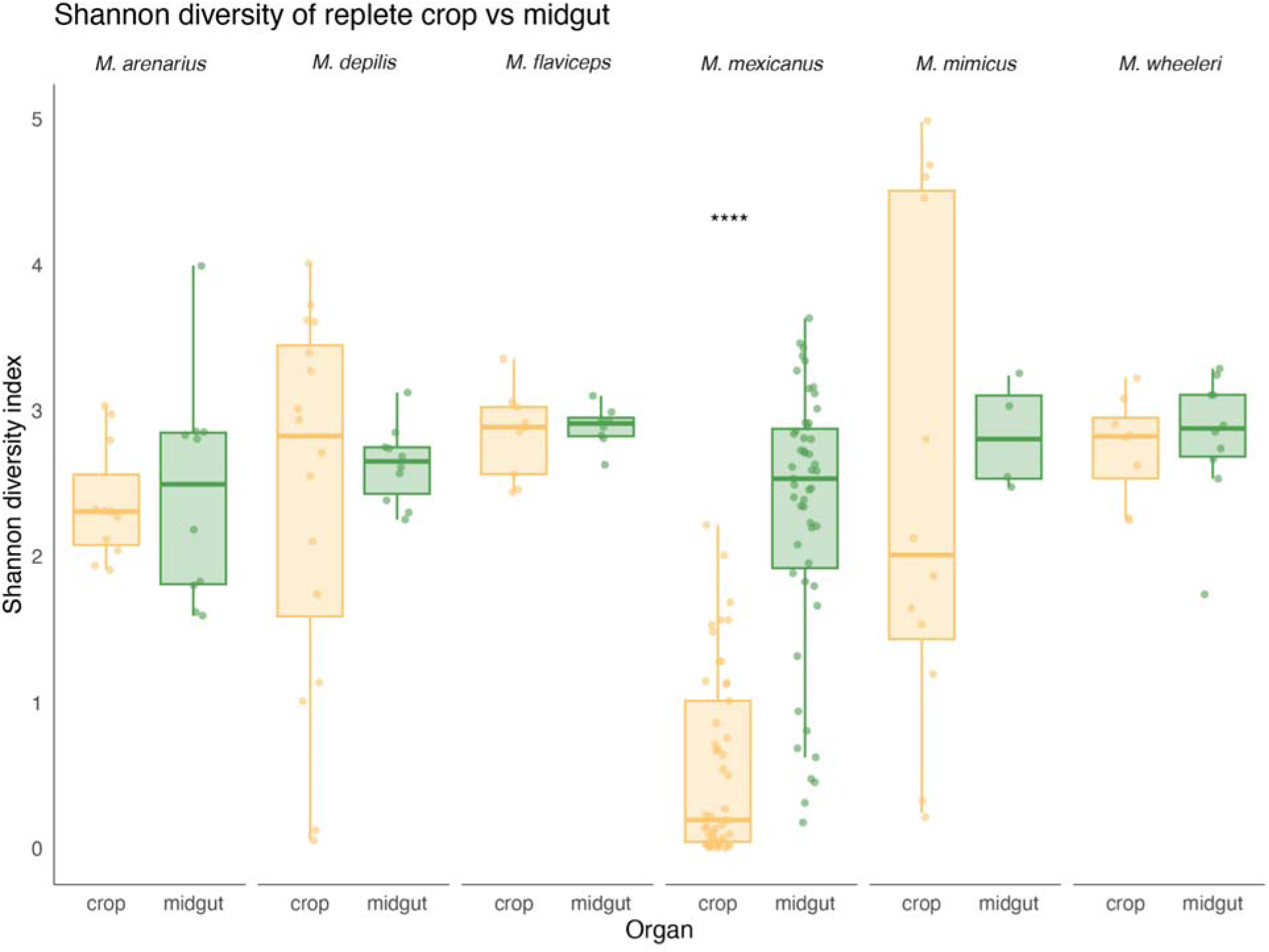
Shannon diversity comparing replete crop and midgut microbiome across six *Myrmecocystus* species, with each point representing individual samples. Sample size (crop/midgut) was *M. arenarius* (11/10), *M. depilis* (16/10), *M. flaviceps* (9/8), *M. mexicanus* (49/51), *M.* mimicus (12/4), *and M. wheeleri* (8/10). The midgut communities generally exhibit higher diversity than crop communities, however, this difference was statistically significant only in *M. mexicanus* (Wilcoxon rank-sum test, FDR-adjusted p = 9.55e^-13^). No significant differences in Shannon diversity between the crop and midgut samples were observed in other *Myrmecocystus* species (all FDR-adjusted p > 0.05).

To further examine the bacterial taxa associated with these organ differences, we used ANCOM-BC2 to compare unrarefied sequence counts between *M. mexicanus* replete crop (n = 52) and midgut (n =52). We identified several genera that differed significantly in abundance between the crop and midgut of *M. mexicanus* repletes (Fig. 5; Table S3). *Fructilactobacillus* and Acetobacteraceae G004843345 were more abundant in the crop, whereas *Uruburella, Streptococcus, Pseudoalteromonas,* and *Vibrio A* 676397 were differentially more abundant in the midgut (Fig. 5; Table S3). We found no differentially abundant taxa between the crop and midgut in *M. depilis*, *M. mimicus, M. arenarius, M. flaviceps,* and *M. wheeleri*.

**Figure 5.**
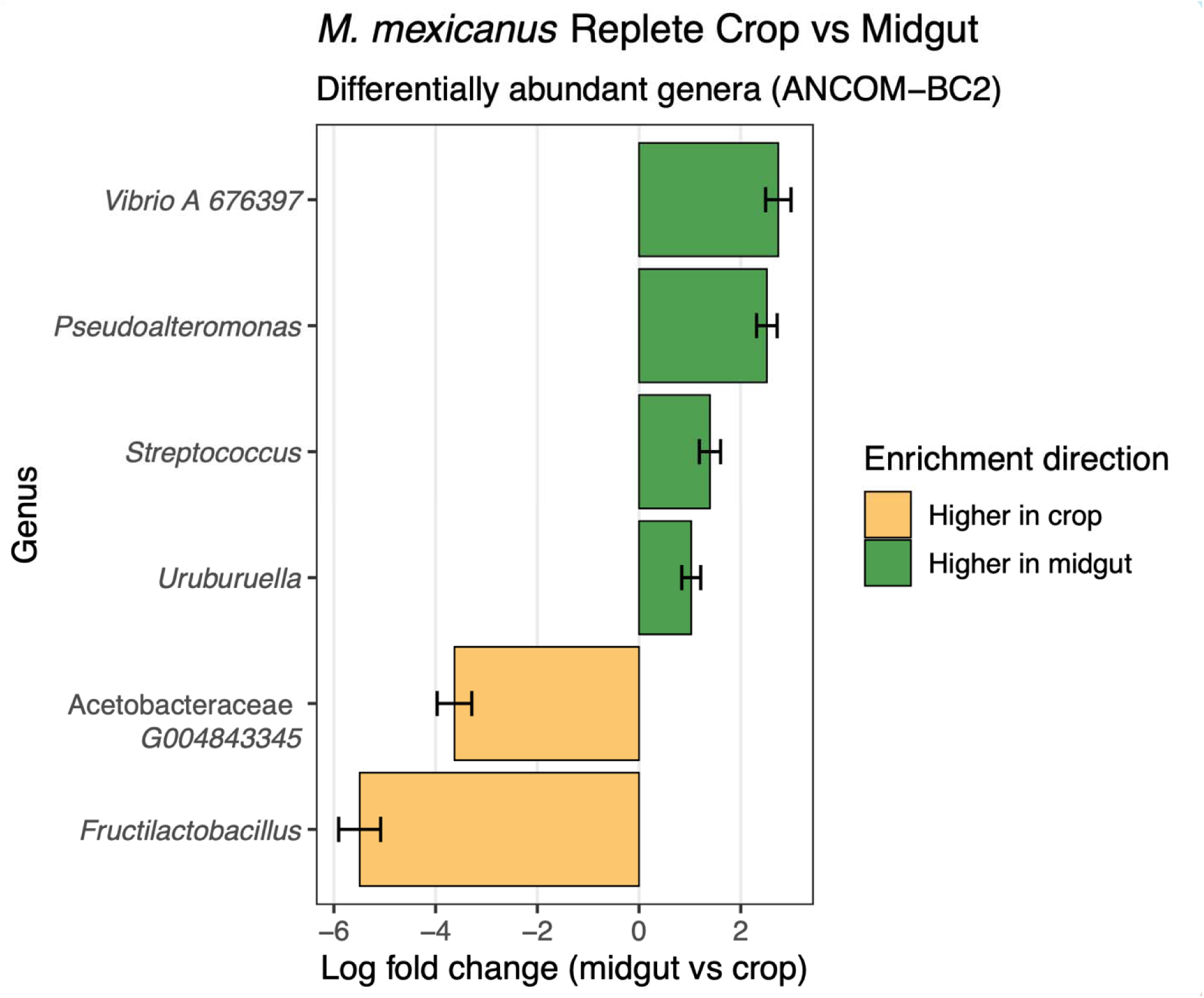
Differentially abundant bacterial genera between 104 *M. mexicanus* replete crop (n = 52) and midgut (n = 52) samples identified using ANCOM-BC2. Negative Log_2_ fold changes (yellow) indicate genera differentially abundant in the crop, while positive log_2_ fold change values (green) indicate enrichment in the midgut. *Fructilactobacillus* and Acetobacteraceae G004843345 were more significantly enriched in *M. mexicanus* crop, whereas *Uruburella, Streptococcus, Pseudoalteromonas,* and *Vibrio A 676397* were significantly enriched *M. mexicanus* midgut. The error bars represent standard errors of the estimated log_2_ fold changes.

Overall, our results suggest that gut compartmentalization varies among *Myrmecocystus* species. In particular, *M. mexicanus* showed the clearest compartmentalization pattern, with significant differences in crop and midgut microbiome composition, alpha diversity, and the abundance of specific bacterial taxa.

## Discussion

In this study, we characterized gut microbiomes across six *Myrmecocystus* species by analyzing the communities across species, colony, caste, and organ. Across our analyses, microbiome composition was primarily driven by ant species identity. Caste had a limited overall influence on microbiome structure, with repletes and workers generally retaining similar bacterial communities despite differences in morphology, varying tasks, and potential environmental exposures. While at first we were surprised that there were minimal differences between castes, in retrospect it makes sense, since these ant species engage in frequent trophallaxis, constantly sharing the contents of their digestive systems (18). Organ-level differences between gut microbiomes were statistically significant in the species *M. mexicanus, M. wheeleri,* and *M. mimicus*. While colony-level differences in gut microbiome composition were statistically significant for *M. mexicanus, M. arenarius, and M. depilis,* they were not consistent across all species. Among all the species examined, *M. mexicanus* showed evidence for a resident gut microbiome, with replete crops dominated almost entirely by *Fructilactobacillus*, across four years of sampling and eight colonies. This monodominance is striking when compared to other species within the *Myrmecocystus* genus, which generally retained more diverse crop microbial communities.

Gut compartmentalization can play a role in shaping the microbial community. For example, in *Cephalotes* turtle ants, specific gut compartments support specialized and conserved symbiont communities (5). Here, we observe evidence of compartmentalization in *M. mexicanus:* the crop was dominated by *Fructilactobacillus*, while the midgut showed greater microbial diversity. While *M. mexicanus* has a striking resident microbiome and the other species have more diverse resident microbiomes, the results of this study do not tell us if the gut microbiomes serve any function for the ant, or if the bacteria are merely commensals living in an environment that is suited for them. However, based on the identities of the community members, we can form testable hypotheses for future work in determining the functional roles of these microbes.

### Lactic acid and acetic acid bacteria

The complete dominance of *Fructilactobacillus* was only observed in *M. mexicanus* replete crops, but it was also abundant in other species, just to a lesser degree (e.g., *M. depilis*). Where there was less *Fructilactobacillus,* there were often other genera of lactic acid or acetic acid bacteria present in high abundance (Fig. 3). Lactic acid bacteria, particularly, are often found in the guts of sugar-feeding social insects. In aphid-tending *Formicine* ants and *Lasius* ants, their infrabuccal pockets and crops are abundant in lactic acid bacteria such as *Lactobacillus*, *Weissella*, and *Fructobacillus* that are capable of catabolizing sugars commonly found in hemipteran honeydew, including sucrose, trehalose, melezitose, and raffinose (19). Beyond catabolizing sugar-rich foods, lactic acid bacteria can inhibit microbial competitors by producing lactic acid and other antimicrobial compounds, lowering pH, and generating acidic conditions that many bacteria and fungi cannot tolerate, as demonstrated in *Mycocepurus smithii* (20). *Myrmecocystus* ants belong to the subfamily Formicinae, which are known to produce and consume formic acid from modified poison glands. Some ants in this subfamily have been shown to control their gut microbiomes by creating an acidic crop environment through formic acid consumption (21). *Fructilactobacillus*, other lactic acid bacteria, and acetic acid bacteria, which are acid-tolerant, could be abundant due to these conditions in *Myrmecocystus* guts, or they could be contributing to further acidification to control their environment in their favor. *Fructilactobacillus* is widely used in the food industry for the fermentation of dairy, meat, sourdough, and vegetables (20, 22–24). Acetic acid (vinegar) is also used in food production and preservation, both with live culture fermentation and with industrially derived vinegar (25, 26). Acetic acid bacteria inhibit the growth of foodborne pathogens (e.g., *Salmonella* spp.), as well as agents of food spoilage such as yeasts and molds (27).

The acid-loving bacteria that dominate the digestive systems of *Myrmecocystus* ants could be commensal, not providing any benefit or harm to their host, but simply growing in an ideal niche. However, they could also be providing benefits to their hosts. First, they may facilitate the metabolism of sugar-rich diets by efficiently catabolizing carbohydrates (e.g., hemipteran honeydew and extrafloral nectar) stored in replete crops into sugars or other molecules that are more easily metabolized by the ants. Second, they may contribute to host fitness by acidifying the gut environment to inhibit the growth of pathogenic microbes. And third, their function may be analogous to lacto-fermentation for human food preservation, where *Fructilactobacillus* prevents spoilage of nutrients in replete crops. In future work, we will examine the growth and metabolism of *Myrmecocystus* lactic acid and acetic acid bacteria, starting with *Fructilactobacillus* in *M. mexicanus* guts.

### Halophiles

We also detected halophilic archaea and bacteria, including *Haloparvum, Haloarcula, Halorubrum, Halobacterium,* and *Natronomonas* in multiple *Myrmecocystus* species collected in California. The high osmotic pressure generated by concentrated sugars in replete crops may favor the proliferation of halotolerant taxa, not because they are necessarily adapted to high salt environments, but because they are physiologically adapted to high osmotic pressures. Both high-salt and high-sugar environments reduce water activity and impose similar constraints on microbial growth (28). These halophiles could be adapted to the high-sugar conditions within the crop. In *Myrmecocystus,* replete crops are concentrated in carbohydrates, composed primarily of fructose and glucose, with smaller amounts of maltose (29). Strains of *Tetragenococcus halophilus,* typically associated with high-salt environments, have also been detected in sugar-rich beet juice (30, 31). Justé et al. (2008) found that some strains of *T. halophilus* were adapted to high salt, while others were adapted to high sugar environments, but not both (30). Halophilic archaea, including *Halobacterium* and *Haloarcula,* are capable of growing on sugars such as glucose, fructose, and sucrose (32). Together, these observations raise the possibility that halophilic bacteria and archaea detected in *Myrmecocystus* guts represent lineages adapted to high sugar conditions within the replete crop.

### Where do the microbes come from?

The variation we see in microbiome composition across *Myrmecocystus* spp. is expected in social insects, where species in the same genus may differ in diet, habitat, foraging behavior, and exposure to environmental conditions (33). Even within *M. mexicanus*, variation among colonies and sampling periods suggests that beyond ant species identity, differences in vegetation, resource availability, and soil composition across habitats may influence microbial assembly across colonies. Similar patterns are observed in other social insects, where microbiome composition reflects environmental context. For example, *Oecophylla smaragdina* weaver ant gut microbiome composition differs between forest and urban habitats primarily due to variations in feeding habits and host diet (34).

Diet could contribute to the structuring of the *Myrmecocystus* microbiome. *Myrmecocystus* foragers collect a wide range of resources, including nectar, gall secretions, aphid and coccid secretions, and extrafloral nectar (29). While all *Myrmecocystus* are considered omnivorous scavengers, there are differences in dietary preferences and foraging habits among species (35). Species that are primarily nocturnal, such as *M. mexicanus*, are thought to largely rely on liquid carbohydrate resources such as nectar and plant exudates (11, 35). Whereas more diurnal species, such as *M. wheeleri,* are described as more carnivorous, supplementing their diet with live or dead arthropods (11, 35). Comparative studies on Indo-Pacific ants identified the host’s dietary niche as the primary driver of microbial diversity and outweighed the physical environment (36). In many cases, microorganisms do not just occupy the crop but actively define its physicochemical state. In *Apis mellifera* honeybees, their diet, consisting of nectar and honey, introduces high levels of sucrose, glucose, and fructose in the crop, acting as a selective pressure that controls the assembly of local bacterial and fungal communities (37). Microorganisms like acetic acid bacteria act as oxygen scavengers to maintain subtoxic conditions, while other fermentative bacteria produce organic acids that acidify the gut environment (37). We suspect that a similar process is occurring in *Myrmecocystus,* where their diet favors microbes adapted to carbohydrate metabolism, low pH tolerance, and osmotically stressful environments in the crop. We expect that many members of the *Myrmecocystus* gut microbiome originate from their diets. Many lactic acid bacteria that have been found in the crops of these ants have also been isolated from nectar (38). It is also likely that halophilic archaea and bacteria are also found in sources such as nectar with relatively high osmotic pressure. In the human gut, it has been found that halophilic archaea have been associated with the dietary intake of fermented and salted foods (39, 40). We expect that diet could be a source for microbiome community members, and that the crop conditions help to further shape the community.

Another possible contributor to the origin of *Myrmecocystus* gut microbiome is through contact with their immediate surroundings, such as soil-associated microbes. From our collected samples, many of the colony entrances were located in close geographic proximity, which could increase the likelihood of shared environmental exposure. In *Veromessor andrei* harvester ants, colonies that are geographically closer harbor more similar microbial communities (41). However, microbiome composition in this system is also shaped by host-associated factors such as diet, life stage, and caste (41). If soil-associated microbes were a dominant driver of microbiome assembly in *Myrmecocystus,* we would expect colonies that are spatially close, regardless of species identity, to have similar microbial communities. Yet our results showed that colonies, such as *M. mexicanus* (X9 and X10) and *M. mimicus* (M1 and M2) collected in July 2025, retained distinct species-specific microbiome compositions despite being in close proximity (Fig. 3). Therefore, we do not think that the immediate environment, namely the soil, is the main contributor to the microbiome of these ants.

### Moving forward

The results presented here lay the groundwork for many future studies. We will work to disentangle the ecological and evolutionary drivers shaping these microbial communities. At the core, we want to answer the question: do honeypot ant microbiomes play a role in allowing them to maintain such extreme life histories?

## Materials and Methods

### Field Sampling

*Myrmecocystus* ants were collected at five time points from 2022-2025 (Table S1), either from private property with permission or Bureau of Land Management (BLM) land, throughout the Southwestern United States (Table S1). Initial species identification was conducted onsite using worker morphological characteristics. We later confirmed ant identification through sequencing and analysis of the COI gene (11). Overall, we collected six *Myrmecocystus* species including *M. arenarius, M. depilis, M. flaviceps, M. mexicanus, M. mimicus*, and *M. wheeleri*. Our goal was to collect 10 repletes per colony, although this was not always achievable (Table S1). Alongside replete and worker samples, we also collected queens, alates, and brood; however, these collections were inconsistent and are saved for future examinations.

To collect repletes and workers, we excavated colonies until we found chambers containing repletes. Compared to repletes, regular workers were easy to find and were located both in the replete chambers and throughout the rest of the colony chambers and tunnels. Typically, replete chambers would be found at least 1 m underground, but on rare occasions we would find them closer to the surface. Holes were carefully hand-dug with small shovels and were filled in once collections were completed. We removed repletes and workers with featherweight forceps or with fan paintbrushes into containers. The ants were kept in plastic containers with a moistened paper towel for 24 to 48 hours. They were not fed and were kept alive until the dissection process. Ants from particular colony fragments were maintained together prior to dissection, allowing possible trophallactic exchange to continue during this period. Ants were first surface sterilized with 70% ethanol followed by sterile phosphate-buffered saline. Dissections were carried out under sterile conditions next to an open flame, using a dissecting scope, on a sterile petri dish. To minimize microbial contamination, all tools and equipment were surface sterilized repeatedly between each dissection using 70% ethanol and by passing them through the flame. The ants were surface sterilized and dissected by first removing the head, then the external membrane of the gaster was opened using sterile ultrafine forceps, and the digestive organs (crop and midgut) were transferred with contents intact and not leaking into individual sterile tubes. If any organ ruptured during dissection, the sample was discarded and not used for DNA extraction or subsequent analyses. Dissected midguts and crops were placed into tubes containing 100 μL of TES buffer (10mM pH 7.5 Tris, 10mM pH 8 EDTA, 0.5% SDS) for DNA extractions. For the repletes, dissected midguts were also placed into 100 μL of TES buffer. Because the replete crops contained a relatively large amount of fluid, we reserved some material for future proteomics, metabolomics, and microbial isolation. However, this study is focused on characterizing the microbiome community composition, and so we describe only the methods for conducting community amplicon sequencing. We included negative controls with every batch of DNA extractions. We used blank tubes with the same TES buffer as the samples and performed all DNA extraction, library prep, and sequencing steps alongside the samples. We had at least two negative controls included for each collection trip and sequencing run, and these are available in the raw data files.

### DNA Extractions and Sequencing

For individual ant crop and midgut samples stored in 100 μL of TES buffer (10mM pH 7.5 Tris, 10mM pH 8 EDTA, 0.5% SDS), our custom phenol/chloroform extraction protocol was followed. Samples were incubated with 36 μL of lysozyme (100 mg/mL in 10mM Tris pH 7.5; Thermo Scientific, Waltham, MA, USA) for 24 hours at 37 . Then, 220 μL of TES and 5 μL proteinase K (800 U/ml; New England Biolabs, Ipswich, MA, USA) were added to our samples. Samples were incubated at 56°C for 20 minutes. Following incubation, 250 μL of Sigma Aldrich 3M sodium acetate was added, and samples were frozen at -80°C for at least 15 minutes. Once the samples had thawed, samples were spun at 14,000 rpm for 15 minutes at 4°C. The supernatant is then transferred to a new tube, and a 1:1 volume of phenol:chloroform is added. Samples were spun at 14,000 rpm for 15 minutes at 4 . The supernatant was transferred to a new tube, and a chloroform wash was performed. Before precipitating, 10% sodium acetate and 90% cold isopropanol were added. Samples were spun at 14,000 rpm for 15 minutes at 4°C to pellet the DNA. Then we performed two ethanol washes with 70% ethanol. The DNA was then resuspended in 50 μL TE (10mM pH 7.5 Tris, 1mM EDTA). DNA concentrations were quantified with Qubit HS dsDNA (Thermo Fisher Scientific, Waltham, MA, USA). Extracted DNA samples were then shipped to the University of Connecticut’s Microbial Analysis, Resources, and Services (UCONN MARS) facility for 16S rRNA gene amplicon sequencing and ITS amplicon sequencing.

According to the UCONN MARS facility protocol, the 16S rRNA V4 region was amplified using 515F and 806R with Illumina adapters and dual 8-basepair indices (42). The ITS2 region was amplified with ITS3 and ITS4 primers (43) using the same dual indexing design as the V4. PCR reactions were performed with an initial denaturation of 95°C for 3.5 minutes, followed by 30 cycles of 30 seconds at 95.0°C, 30 seconds at 50.0°C, and 90 seconds at 72.0°C, with a final extension at 72.0°C for 10 minutes. PCR products were pooled for quantification and visualization using the QIAxcel DNA Fast Analysis (Qiagen, Hilden, Germany). PCR products were normalized based on the concentration of DNA from 250-400 bp, then pooled using the epMotion 3075 liquid handling robot. The pooled PCR products were cleaned using Mag-Bind Beads (0.8x ratio; Omega Bio-Tek, Norcross, GA, USA) and sequenced on an Illumina MiSeq platform (Illumina; San Diego, CA, USA) with a v2 2×250 base pair kit, for paired-end sequencing. We only conducted ITS sequencing for the first set of collected samples. Because we found almost no fungi in our samples (Fig S6-S7), we chose to focus on bacteria and archaea only in subsequent sequencing runs.

### Amplicon and Statistical Analysis

We processed, aligned, and categorized sequence reads using DADA2 1.25.2 in R (44). To do so, we first created read quality profiles to visually inspect before filtering and trimming. Forward and reverse reads were truncated to 240 and 160 base pairs. While any reads containing ambiguous bases, exceeding two expected errors in either direction, or matching PhiX were removed. Our filtering parameters also included a minimum read length of 10 bp and a quality score truncation threshold of 2. Error rates were independently estimated for forward and reverse reads, followed by denoising and amplicon sequence variant (ASV) inference using dada (). Paired-end reads were merged using mergePairs (), and chimeric sequences were identified and removed using removeBimeraDenoco() with the consensus method. We then used the prevalence method, with a threshold of 0.1, in decontam to distinguish potential contaminants from true biological sequences found in our negative controls (45).

After filtering, trimming, merging reads, and removing contaminants, amplicon sequence variants (ASV) were taxonomically assigned using the Greengenes v2 database. As a note, we did initially run our analysis using the SILVA database but switched to Greengenes because it provided more successful matches to our data. To construct the ASV-based phylogeny required for weighted UniFrac analyses, we first made an alignment using MAFFT (Katoh et al., 2002). An ASV phylogeny was inferred from these sequences using the GTR+G model in IQtree v2.4.0 (46). The resulting tree was rooted in R using the phangorn package (47). The rooted tree was then incorporated into our phyloseq object (48). Rarefaction curves were then generated to assess sequence depth (Fig. S1). Samples containing fewer than 3,000 reads were excluded, and the remaining samples were rarefied to 3,000 reads per sample for downstream diversity analyses.

Weighted UniFrac distances were calculated from the decontaminated, rarefied dataset using the phyloseq package, and ordinations were performed using non-metric multidimensional scaling (NMDS) with the vegan package (49). Microbiome composition was visualized using NMDS plots generated with the plot_ordinate function in phyloseq (48).

All data analysis was run using decontaminated sequencing data. For some tests, it was recommended that rarefied data be used (samples containing fewer than 3,000 read counts were excluded), while other tests required unrarefied data (included all reads). Statistical significance between groups was evaluated using global permutational multivariate analysis of variance (PERMANOVA) and pairwise differences with analysis of similarity (ANOSIM) in the vegan package (49). To further investigate the extent of clustering across species, colony, caste, and organ, we performed permutational multivariate analysis of dispersion (PERMDISP) to measure beta diversity (50). Analysis of compositions of microbiomes with bias correction (ANCOM-BC2) was used to identify bacterial genera differentially abundant between the crop and midgut of replete workers within each *Myrmecocystus* species (51). ANCOM-BC2 was primarily used for organ comparison because it allowed us to test whether the repletism phenotype is associated with differential abundance of specific taxa. ANCOM-BC2 analyses were performed on unrarefied data at the genus level, with organ set as the fixed effect. The crop was used as the reference group; therefore, negative log fold changes indicated greater abundance in the crop and positive values indicated greater abundance in the midgut. Genera that were detected in fewer than 10% of samples and samples with 1,000 reads were excluded from the analysis. Structural zero-inflation and pseudo-count sensitivity analyses were used. P-values were adjusted for multiple testing using the Holm method, with significance set to 0.05. Alpha diversity metrics, including observed richness, Shannon diversity, and inverse Simpson diversity, were calculated from our rarefied dataset (321 samples) using phyloseq to measure the richness and evenness of microbial communities within the samples (48, 52, 53). Statistical differences in the Shannon diversity index were first assessed using the Kruskal-Wallis test, followed by pairwise Wilcoxon rank-sum tests with Benjamini-Hochberg correction for multiple comparisons (54–56). Afterwards, we used a Gaussian generalized linear mixed model (GLMM) with the glmmTMB package (57) and non-rarefied data (404 samples) to compare the Shannon diversity patterns found with our Kruskal-Wallis and Wilcoxon tests. We included species, organ, their interaction, and caste as fixed effects. Colony and individual ant identity were included as random effects to account for colony-level variation and paired crop and midgut samples. Post hoc pairwise comparisons were performed using estimated marginal means with the emmans package. Bubble plots were generated in R using the ggplot2 and reshape2 packages to visualize the presence and relative abundance of the top 50 most abundant microbial taxa across species and across organs (58, 59).

## Data Availability

All raw sequencing data are available in the NCBI Sequence Read Archive (SRA) under the Bioproject PRJNA1449520: *Myrmecocystus* honeypot ants 16s rRNA. Metadata and code used for analysis are available on GitHub at: https://github.com/dqvnguyen/HPA-Microbiome.

## Acknowledgements

The project was funded by NSF BRC-BIO grant 2312984 and seed funding through Rutgers Global to LK and an NSF PRFB Award 2305685 to CF.

## Author Contributions

We followed CRediT’s 14 Contributor Roles guidelines. **DVN**: Conceptualization; Data Curation; Formal Analysis; Investigation; Validation; Visualization; Writing (original draft); Writing (review and editing). **CF**: Conceptualization; Data Curation; Funding Acquisition; Investigation; Methodology; Visualization; Writing (original draft). **BN**: Investigation. **IS**: Investigation; Methodology. **ML**: Investigation; Methodology. **LK**: Conceptualization; Funding Acquisition; Project Administration; Supervision; Validation; Writing (review and editing)

## Notes

### Competing Interest Statement

The authors have declared no competing interest.

### Summary of Updates

This version of the manuscript has been revised to be more specific about our materials and methods, providing details on our 16S rRNA gene sequencing procedure, quality control, negative controls included during our DNA extraction process, and bioinformatic analysis. Additionally, the revised manuscript incorporates statistical approaches beyond PERMANOVA, ANOSIM, and ANCOMBC to better understand community clustering and evaluate differences in microbial community composition and diversity, such as PERMDISP and generalized linear mixed models. This version contains our revised figures and supplemental materials, such as our stacked bar charts, ITS sequencing, and tables that include sample size to improve clarity.

