## Supplemental Figures for "*Myrmecocystus* honeypot ants have a species-specific resident gut microbiome"

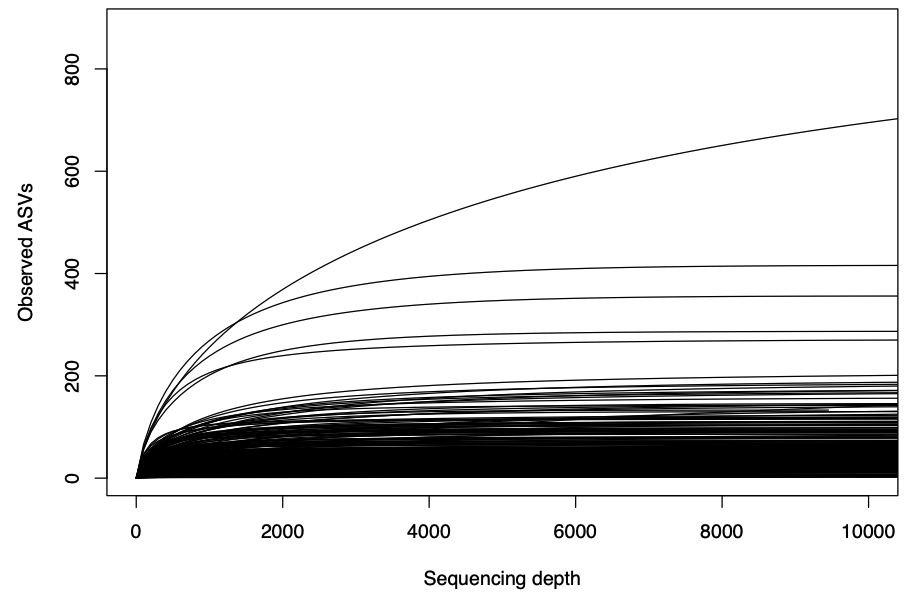

Figure S1. Rarefaction curves showing the relationship between sequencing depth (number of reads) and observed amplicon sequence variant (ASV) across samples. Each line represents an individual sample. Most curves approached a plateau at relatively low sequencing depth of approximately 3,000–5,000 reads. Based on these curves and sample retention (78.5%; 324 samples), 3,000 reads per sample was selected as the rarefaction depth for downstream diversity analyses.

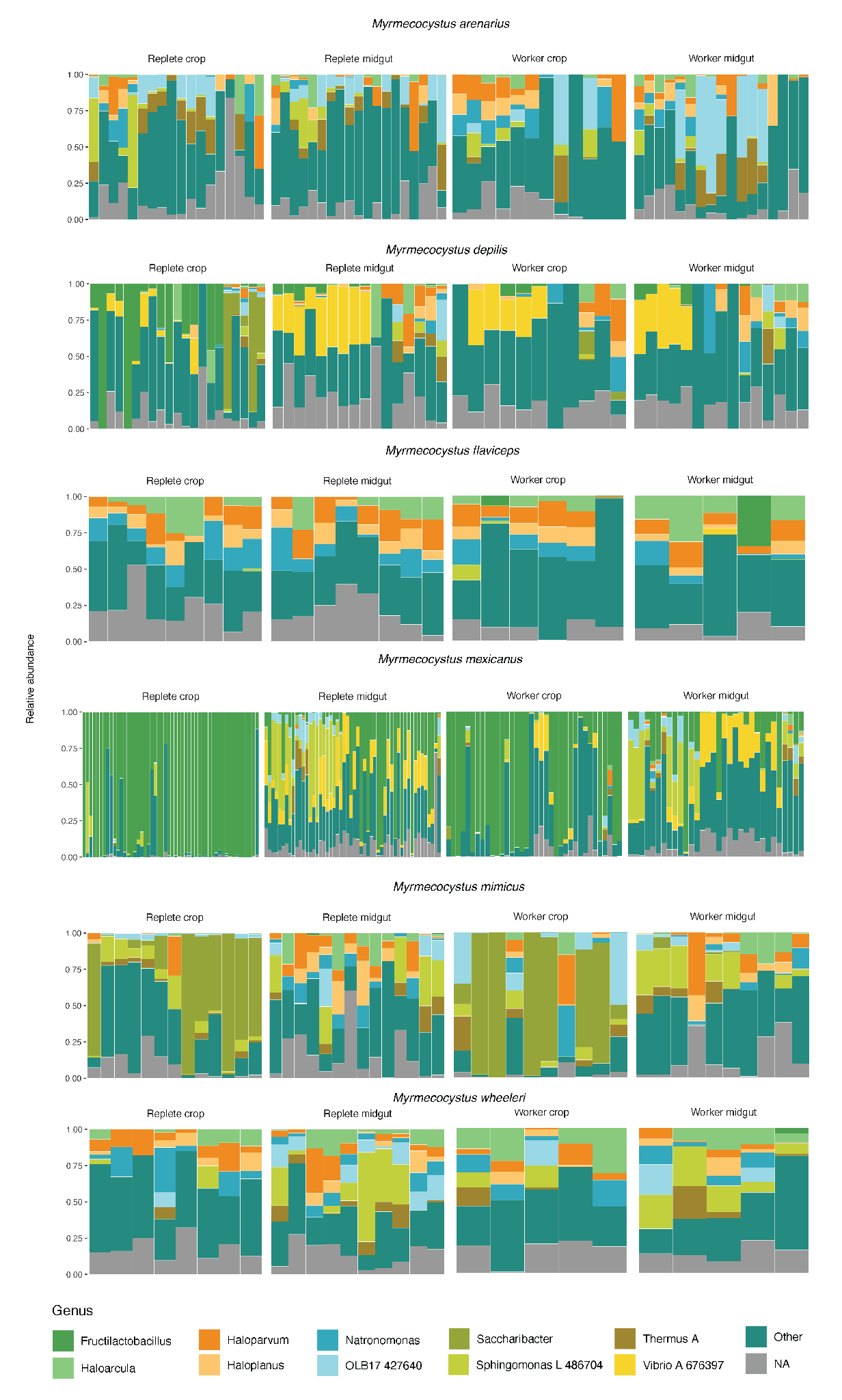

Figure S2. Relative abundance of bacterial composition across six *Myrmecocystus* species*: M. arenarius* (n = 35)*, M. depilis* (n = 39)*, M. flaviceps* (n = 27)*, M. mexicanus* (n = 166)*, M. mimicus* (n = 26)*,* and *M. wheeleri* (n = 28)*.* Within each species, samples are grouped by caste and organ (replete crop, replete midgut, worker crop, and worker midgut). Each bar represents an individual sample, and colors indicate the 10 most abundant bacterial genera identified across the full dataset, the remaining genera are grouped as “other” and unclassified genera are shown as “NA.” Community composition varies among species, with *M. mexicanus* crop samples showing dominance by a few genera, *Fructilactobacillus,* whereas other species display more diverse and variable community structure.

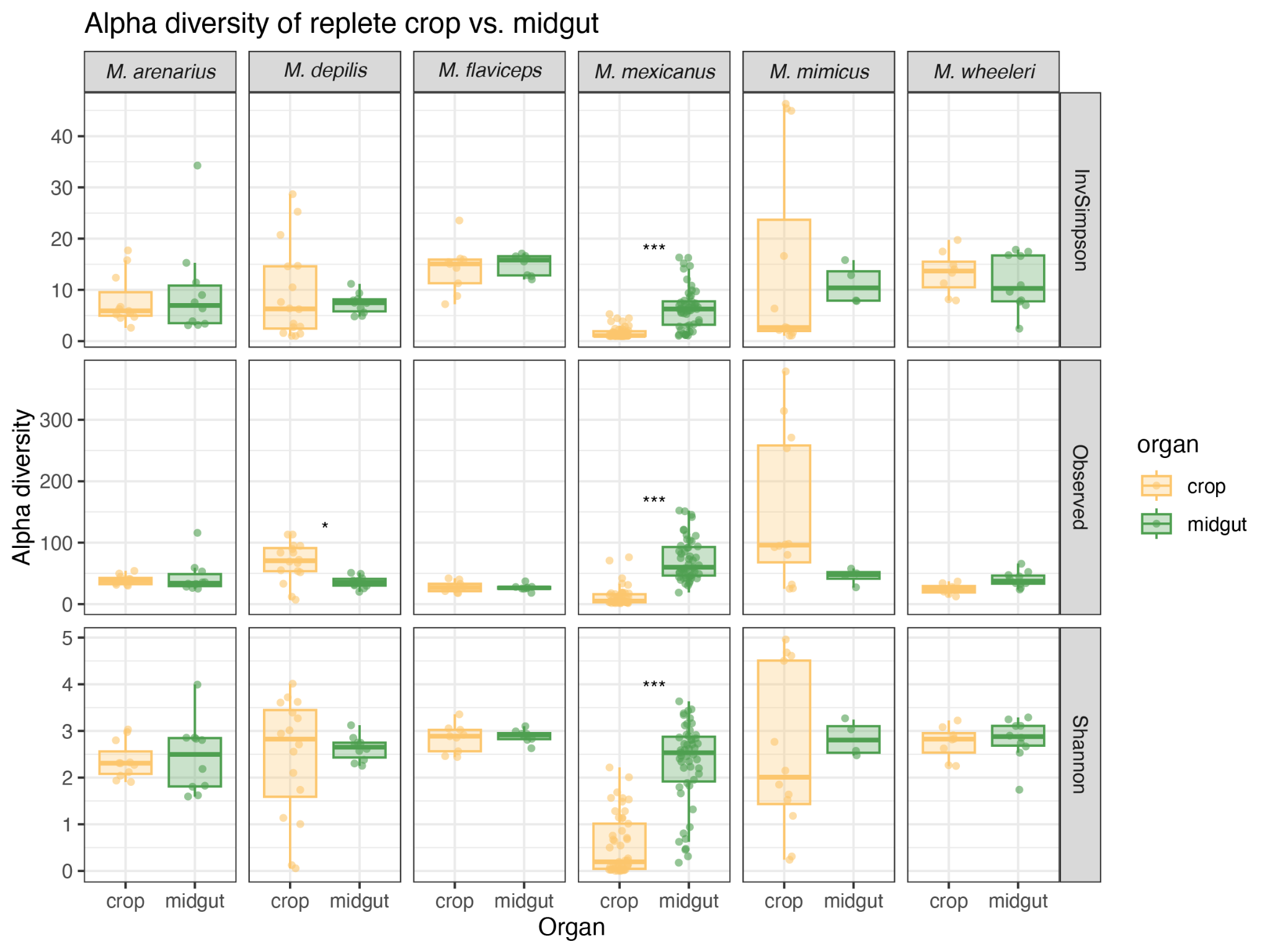

Figure S3. Alpha diversity comparisons between crop and midgut samples across *Myrmecocystus* species. The boxplots show three diversity metrics (Inverse Simpson, Observed richness, and Shannon), with each point representing individual samples. Sample sizes (crop/midgut) were *M. arenarius* (11/10), *M. depilis* (16/10), *M.flaviceps* (9/8), *M. mexicanus* (49/51), *M. mimicus* (12/4), *and M. wheeleri* (8/10). Crop and midgut samples are shown within each species panel. In *M. mexicanus,* the midgut samples exhibited significantly higher alpha diversity than crop samples across all three metrics (Wilcoxon rank-sum test, FDR-adjusted p < 0.001). There were no significant crop and midgut differences detected for the other *Myrmecocystus* species after FDR correction.

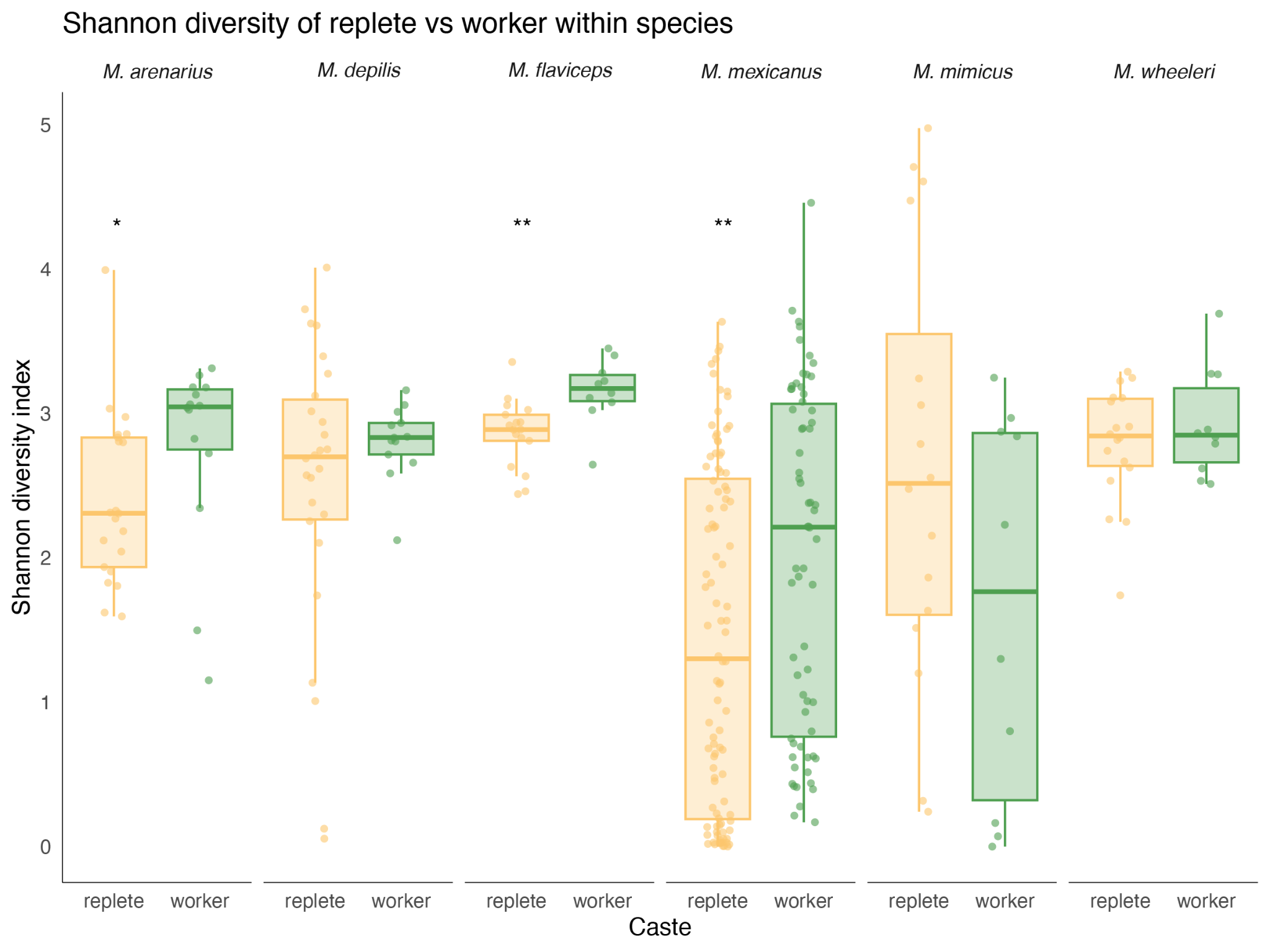
Figure S4. Shannon diversity metric used to compare microbial diversity between replete and castes across six *Myrmecocystus* species. The boxplots show the distribution of Shannon diversity values for replete (yellow) and worker (green) within each species, with each point representing individual samples. The sample size (replete/worker) were *M. arenarius* (21/14), *M. depilis* (26/13), *M. flaviceps* (17/10), *M. mexicanus* (100/66), *M. mimicus* (16/10), and *M. wheeleri* (18/10). Workers exhibited significantly higher Shannon diversity than repletes in M. arenarius, M. flaviceps, and M. mexicanus. However, in other species, such as *M. depilis, M. mimicus,* or *M. wheeleri* there was no significant difference between castes. Significance was determined using Wilcoxon rank sum tests with BH false discovery rate correction (* adjusted p < 0.005; ** adjusted p < 0.01).

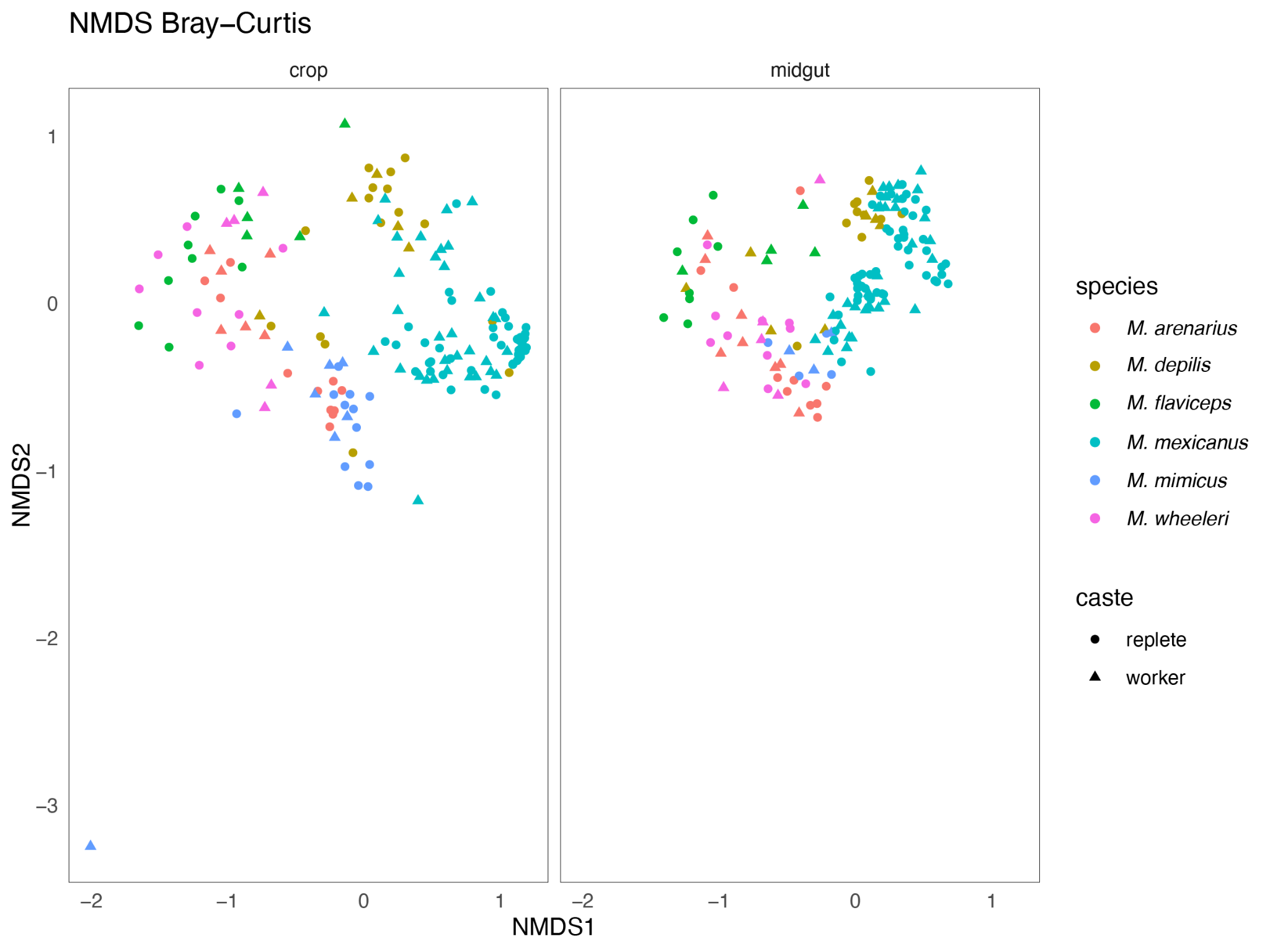

Figure S5. Non-metric Multidimensional Scaling (NMDS) ordination based the Bray Curtis dissimilarity matrix to visualize differences in microbial community composition across *Myrmecocystus* species, caste, and organ. Each point represents individual samples, species indicated by color, caste by shape, with panels corresponding to crop or midgut samples. The ordination includes 321 organ samples across the six species: *M. arenarius* ( n = 35), *M. depilis* (n = 39), *M. flaviceps* (n = 27), *M. mexicanus* (n = 166), *M. mimicus* (n = 26), and *M. wheeleri* (n = 28). Samples show clear species-associated clustering with partial overlap among species. The overall pattern was broadly consistent with the weighted UniFrac ordination used throughout the study.

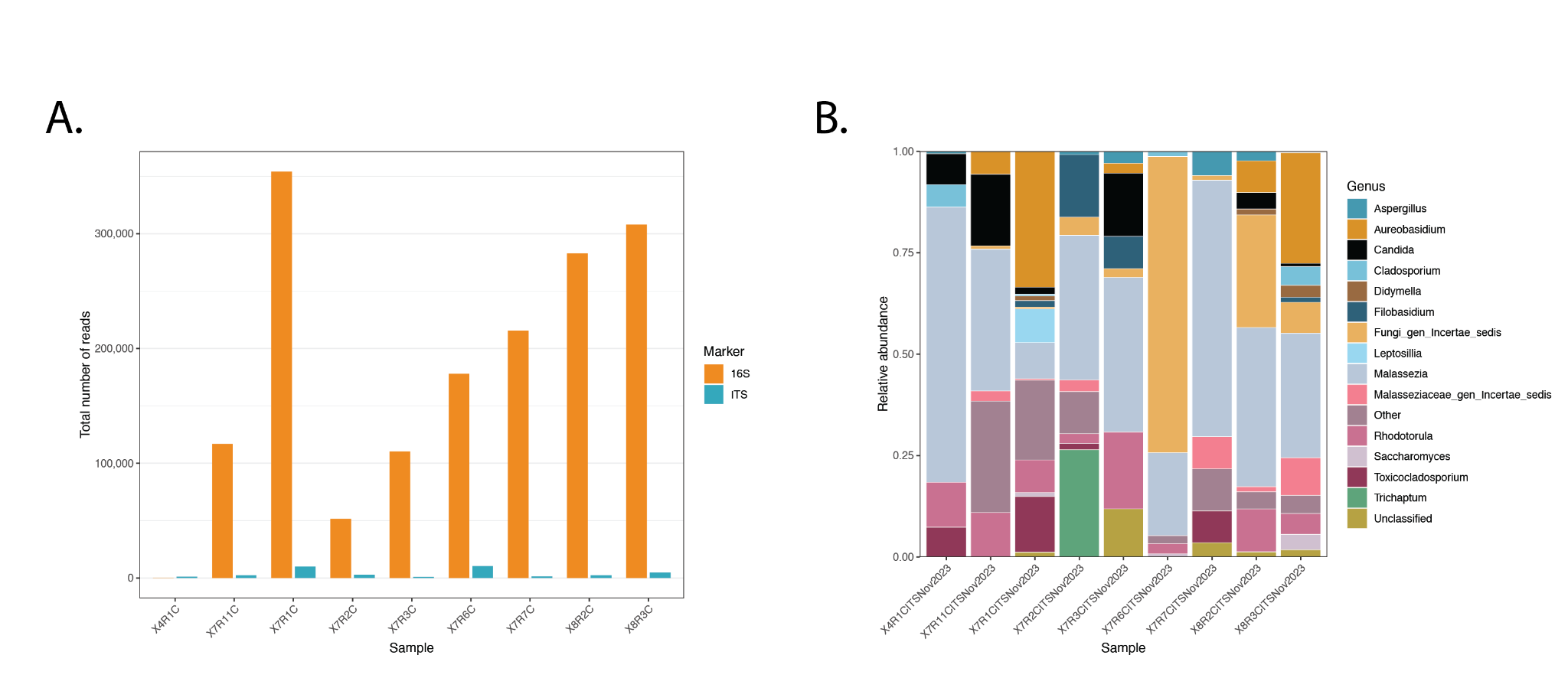

Figure S6. (A) The sequencing depth of 16S and ITS samples from nine *M. mexicanus* replete crops. When placing 16S rRNA gene and ITS amplicon sequencing side by side, we see that ITS samples generally had substantially fewer reads than the corresponding 16S samples. (B) Relative abundance of fungal genera detected in nine *M. mexicanus* replete crop samples using ITS amplicon sequencing. Each bar represents an individual sample, and colors indicate the 15 most abundant genera across samples. This extremely low abundance of fungal reads may be biologically important. We suspect that the bacterial community in the ant guts is excluding fungal growth, thereby preventing CO2 fermentation that could create disruptive bubbles in the ant crops. This is a point for future investigation.
