## Supplemental Tables for "*Myrmecocystus* honeypot ants have a species-specific resident gut microbiome"

Table S1. Myrmecocystus colony collection locations. Collection sites in Arizona and New Mexico are geographically very close to one another. The table also includes the cytochrome c oxidase subunit 1 sequence (COI) used to identify species

| **Colony** | **Year collected** | **Species** | **Repletes** | **Foragers** | **Latitude** | **Longitude** | **State** | **COI NCBI Acession Number** |
| --- | --- | --- | --- | --- | --- | --- | --- | --- |
| A1 | Nov 2024 | *M. arenarius* | 5 | 6 | 33.907671 | -116.668247 | CA | SUB16542139 A1 QB143795 |
| A2 | May 2025 | *M. arenarius* | 3 | 3 | 33.908692 | -116.676865 | CA | SUB16542139 A2 QB143796 |
| A3 | May 2025 | *M. arenarius* | 5 | 3 | 33.90865 | -116.676663 | CA | SUB16542139 A3 QB143797 |
| A4 | May 2025 | *M. arenarius* | 5 | 3 | 33.908946 | -116.677058 | CA | SUB16542139 A4 QB143798 |
| A5 | May 2025 | *M. arenarius* | 3 | 3 | 33.909255 | -116.677404 | CA | SUB16542139 A5 QB143799 |
| D1 | Aug 2023 | *M. depilis* | 10 | 5 | 32.035617 | -109.19375 | AZ | SUB16542139 D1 QB143800 |
| D2 | Aug 2023 | *M. depilis* | 8 | 5 | 32.03595 | -109.194 | AZ | SUB16542139 D2 QB143801 |
| D3 | July 2025 | *M. depilis* | 5 | 6 | 32.094808 | -108.968703 | NM | SUB16542139 D3 QB143802 |
| M1 | July 2025 | *M. mimicus* | 10 | 5 | 32.095799 | -108.972622 | NM | SUB16542139 M1 QB143803 |
| M2 | July 2025 | *M. mimicus* | 5 | 5 | 32.09455 | -108.968594 | NM | SUB16542139 M2 QB143804 |
| V1 | Nov 2024 | *M. flaviceps* | 8 | 6 | 34.450771 | -117.704869 | CA | SUB16542139 V1 QB143805 |
| V2 | Nov 2024 | *M. flaviceps* | 1 | 4 | 34.453058 | -117.705979 | CA | SUB16542139 V2 QB143806 |
| W1 | Nov 2024 | *M. wheeleri* | 6 | 0 | 33.909031 | -116.669478 | CA | SUB16542139 W1 QB143807 |
| W2 | Nov 2024 | *M. wheeleri* | 4 | 5 | 33.9084983 | -116.669716 | CA | SUB16542139 W2 QB143808 |
| X1 | July 2022 | *M. mexicanus* | 10 | 6 | 31.908 | -109.152 | AZ | Morphologically identified |
| X2 | July 2022 | *M. mexicanus* | 6 | 5 | 31.902 | -109.109 | AZ | Morphologically identified |
| X3 | July 2022 | *M. mexicanus* | 5 | 5 | 31.902 | -109.109 | AZ | Morphologically identified |
| X4 | Aug 2023 | *M. mexicanus* | 5 | 5 | 32.035617 | -109.19375 | AZ | Morphologically identified |
| X7 | Aug 2023 | *M. mexicanus* | 19 | 5 | 32.0356333 | -109.1945333 | AZ | Morphologically identified |
| X8 | Aug 2023 | *M. mexicanus* | 4 | 5 | 32.035083 | -109.193433 | AZ | Morphologically identified |
| X9 | July 2025 | *M. mexicanus* | 1 | 5 | 32.09422 | -108.968719 | NM | SUB16542139 X9 QB143809 |
| X10 | July 2025 | *M. mexicanus* | 6 | 5 | 32.094684 | -108.96855 | NM | SUB16542139 X10 QB143810 |

Table S2. Pairwise comparisons of Shannon diversity across Myrmecocystus species using Wilcoxon rank sum tests with continuity correction. Values represent Benjamini Hochberg false discovery rate (FDR)-adjusted p-values for differences in Shannon diversity between species pairs. Adjusted p-values < 0.05 indicate significant differences in Shannon diversity between species.

|  | ***M. arenarius*** | ***M. depilis*** | ***M. flaviceps*** | ***M. mexicanus*** | ***M. mimicus*** |
| --- | --- | --- | --- | --- | --- |
| ***M. depilis*** | 0.58828 | - | - | - | - |
| ***M. flaviceps*** | 0.01417 | 4.70E-02 | - | - | - |
| ***M. mexicanus*** | 0.00059 | 8.70E-05 | 1.50E-06 | - | - |
| ***M. mimicus*** | 0.34277 | 2.80E-01 | 2.80E-02 | 1.40E-01 | - |
| ***M. wheeleri*** | 0.13902 | 3.80E-01 | 2.00E-01 | 1.60E-05 | 0.08898 |

Table S3. ANCOM-BC2 results identify taxa differentially abundant between the crop and midgut in M. mexicanus. Log fold change (LFC) values indicate the direction and magnitude of differential abundance, with negative values corresponding to taxa enriched in the crop and positive values indicating taxa enriched in the midgut. q-values represent Holm adjusted p-values. The differentially abundant column indicates whether a taxon is significantly differentially abundant between organs (q < 0.05).The robustly differential column indicates taxa that remain significant after an additional robustness corrected test. Fructilactobacillus and Acetobacteraceae G004843345 are differentially more abundant in the crop of M. mexicanus than the midgut. While Vibrio, Pseudoalteromonas, Streptococcus, and Uruburuella are differentially more abundant in the midgut than the crop.

| **Taxon** | **LFC** | **Standard error** | **q-value (Holm-adjusted)** | **Differentially abundant** | **Robustly differential** | **Direction** |
| --- | --- | --- | --- | --- | --- | --- |
| Fructilactobacillus | -5.493120909 | 0.4120911833 | 4.75E-22 | TRUE | TRUE | Higher in crop |
| G004843345 | -3.629026732 | 0.3406920079 | 1.05E-13 | TRUE | TRUE | Higher in crop |
| Uruburuella | 1.026038647 | 0.1855623491 | 3.48E-05 | TRUE | TRUE | Higher in midgut |
| Streptococcus | 1.395472931 | 0.2095965577 | 4.01E-07 | TRUE | TRUE | Higher in midgut |
| Pseudoalteromonas | 2.512807539 | 0.2025946375 | 7.65E-16 | TRUE | TRUE | Higher in midgut |
| Vibrio A 676397 | 2.738076729 | 0.2506843779 | 2.08E-14 | TRUE | TRUE | Higher in midgut |

Table S4. Summary of PERMANOVA, ANOSIM, and PERMDISP analyses using weighted UniFrac and Bray-Curtis distance matrices to test the effects of species, colony, caste, and organ on gut microbiome composition in Myrmecocystus. The R2 values are reported for PERMANOVA where it was available. p-values < 0.05 indicate significant differences in microbial community composition. (With weighted UniFrac)

| **Factor** | **Species / Subset** | **Organ** | **Distance Matrix** | **PERMANOVA R²** | **PERMANOVA p-value ANOSIM R** | | **ANOSIM p-value** | **PERMDISP F** |  | **PERMDISP p-value** |
| --- | --- | --- | --- | --- | --- | --- | --- | --- | --- | --- |
| Species, caste, organ | All samples | - | wUnifrac | 0.612 | 0.000999 *** | - | - | 14.4 0.001 *** | | |
| Species, caste, organ | All samples | - | Bray-Curtis | 0.36164 | 0.000999 *** | - | - |  | 6.3169 | 0.001 *** |
| Species | Repletes | Crop | wUnifrac | 0.6994 | 0.000999 *** | 0.8517 | 0.001 |  | 17.925 | 0.001 *** |
| Species | Repletes | Crop | Bray-Curtis | 0.40178 | 0.000999 *** | 0.8776 | 0.001 |  | 19.941 | 0.001 *** |
| Species | Repletes | Midgut | wUnifrac | 0.57702 | 0.000999 *** | 0.6683 | 0.001 |  | 4.9235 | 0.002** |
| Species | Repletes | Midgut | Bray-Curtis | 0.27866 | 0.000999 *** | 0.6311 | 0.001 |  | 3.181 | 0.011 * |
| Colony | *M. arenarius* | Crop | wUnifrac | 0.70163 | 0.000999 *** | 0.7543 | 0.001 |  | 0.5148 | 0.645 |
| Colony | *M. arenarius* | Crop | Bray-Curtis | 0.32246 | 0.000999 *** | 0.4326 | 0.002 |  | 38.213 | 0.001 *** |
|  |  |  |  |  |  | |  | PERMDISP was not evaluated across all colonies because | | |
|  |  |  |  |  |  | |  | A4 and A5 had n = 1; dispersion could only be assessed | | |
| Colony | *M. arenarius* | Midgut | wUnifrac | 0.54679 | 0.003996 ** | 0.3934 | 0.001 | among replicated colonies. | | |
|  |  |  |  |  |  | |  | PERMDISP was not evaluated across all colonies because | | |
|  |  |  |  |  |  | |  | A4 and A5 had n = 1; dispersion could only be assessed | | |
| Colony | *M. arenarius* | Midgut | Bray-Curtis | 0.28294 | 0.1199 | 0.01803 | 0.404 | among replicated colonies. | | |
| Colony | *M. depilis* | Crop | wUnifrac | 0.42045 | 0.000999 *** | 0.7228 | 0.001 |  | 6.1125 | 0.01 ** |
| Colony | *M. depilis* | Crop | Bray-Curtis | 0.28873 | 0.000999 *** | 0.7842 | 0.001 |  | 18.373 | 0.001 *** |
| Colony | *M. depilis* | Midgut | wUnifrac | 0.56337 | 0.000999 *** | 0.9224 | 0.002 |  | 26.142 | 0.001 *** |
| Colony | *M. depilis* | Midgut | Bray-Curtis | 0.37577 | 0.000999 *** | 0.9762 | 0.001 |  | 15.403 | 0.002 ** |
| Colony | *M. mexicanus* | Crop | wUnifrac | 0.22868 | 0.001998 ** | 0.2886 | 0.001 |  | 6.9362 | 0.001*** |
| Colony | *M. mexicanus* | Crop | Bray-Curtis | 0.32044 | 0.000999 *** | 0.4495 | 0.001 |  | 9.6363 | 0.001 *** |
| Colony | *M. mexicanus* | Midgut | wUnifrac | 0.5044 | 0.000999 *** | 0.5137 | 0.001 |  | 0.5912 | 0.758 |

| Colony | *M. mexicanus* | Midgut |  | Bray-Curtis | 0.47037 | 0.000999 *** | 0.5308 | 0.001 | 0.7785 | 0.59 |
| --- | --- | --- | --- | --- | --- | --- | --- | --- | --- | --- |
| Colony | *M. flaviceps* | Crop |  | wUnifrac | 0.06018 | 0.4436 | -0.009804 | 0.463 | 0.3355 | 0.607 |
| Colony | *M. flaviceps* | Crop |  | Bray-Curtis | 0.08538 | 0.1139 | 0.1201 | 0.242 | 0.0742 | 0.796 |
| Colony | *M. flaviceps* | Midgut |  | wUnifrac | 0.00941 | 1 | -0.2955 | 0.846 | PERMDISP was not evaluated because one colony was represented by only one sample (V2, n = 1). | |
| Colony | *M. flaviceps* | Midgut |  | Bray-Curtis | 0.07144 | 1 | -0.4697 | 1 | PERMDISP was not evaluated across all colonies because A4 and A5 had n = 1; dispersion could only be assessed among replicated colonies. | |
| Colony | *M. mimicus* | Crop |  | wUnifrac | 0.09076 | 0.1538 | -0.07882 | 0.757 | 38.744 | 0.001 *** |
| Colony | *M. mimicus* | Crop |  | Bray-Curtis | 0.38377 | 0.000999 *** | 0.5866 | 0.002 | 21.818 | 0.001 *** |
| Colony | *M. mimicus* | Midgut |  | wUnifrac | 0.3337 | 0.17833 | -0.2182 | 0.862 | 4.4034 | 0.094 |
| Colony | *M. mimicus* | Midgut |  | Bray-Curtis | 0.12937 | 0.7223 | -0.3091 | 0.877 | 2.5927 | 0.193 |
| Colony | *M. wheeleri* | Crop |  | wUnifrac | 0.07538 | 0.3996 | 0.1786 | 0.138 | 0.6229 | 0.54 |
| Colony | *M. wheeleri* | Crop |  | Bray-Curtis | 0.07059 | 0.9071 | -0.2745 | 0.956 | 0.6916 | 0.42 |
| Colony | *M. wheeleri* | Midgut |  | wUnifrac | 0.01634 | 0.966 | 0.04067 | 0.287 | 1.4769 | 0.221 |
| Colony | *M. wheeleri* | Midgut |  | Bray-Curtis | 0.0594 | 0.7522 | -0.065 | 0.707 | 0.0121 | 0.915 |
| Caste | All species | Crop |  | wUnifrac | 0.011669 | 0.1419 | 0.04469 | 0.059 | 2.0502 | 0.144 |
| Caste | All species | Crop |  | Bray-Curtis | 0.0129 | 0.03197 * | 0.003617 | 0.406 | 1.1498 | 0.266 |
| Caste | All species | Midgut |  | wUnifrac | 0.00799 | 0.2737 | 0.01973 | 0.134 | 0.9236 | 0.341 |
| Caste | All species | Midgut |  | Bray-Curtis | 0.0063 | 0.4226 | -0.002855 | 0.509 | 1.4992 | 0.244 |
| Caste | *M. arenarius* | Crop |  | wUnifrac | 0.36817 | 0.008991 ** | 0.2281 | 0.044 | 3.8223 | 0.087 |
| Caste | *M. arenarius* | Crop |  | Bray-Curtis | 0.05646 | 0.02498 * | 0.04976 | 0.179 | 0.5505 | 0.445 |
| Caste | *M. arenarius* | Midgut |  | wUnifrac | 0.1544 | 0.09191 | 0.1521 | 0.085 | 0.0479 | 0.826 |
| Caste | *M. arenarius* | Midgut |  | Bray-Curtis | 0.09348 | 0.09091 | 0.1351 | 0.072 | 0.6695 | 0.408 |
| Caste | *M. depilis* | Crop |  | wUnifrac | 0.10816 | 0.06593 | 0.1333 | 0.171 | 0.1512 | 0.712 |
| Caste | *M. depilis* | Crop |  | Bray-Curtis | 0.11234 | 0.02098 * | 0.1547 | 0.085 | 2.0928 | 0.162 |
| Caste | *M. depilis* | Midgut |  | wUnifrac | 0.14889 | 0.095 | 0.1051 | 0.082 | 4.3037 | 0.063 |
| Caste | *M. depilis* | Midgut |  | Bray-Curtis | 0.08398 | 0.144 | 0.09983 | 0.092 | 3.6456 | 0.081 |
| Caste | *M. mexicanus* | Crop |  | wUnifrac | 0.14013 | 0.000999 *** | 0.3308 | 0.001 | 25.506 | 0.001 *** |
| Caste | *M. mexicanus* | Crop |  | Bray-Curtis | 0.08766 | 0.000999 *** | 0.1965 | 0.001 | 11.544 | 0.001 *** |
| Caste | *M. mexicanus* | Midgut |  | wUnifrac | 0.01589 | 0.2557 | 0.05109 | 0.058 | 3.1085 | 0.076 |
| Caste | *M. mexicanus* | Midgut |  | Bray-Curtis | 0.013 | 0.3477 | 0.03269 | 0.129 | 3.2632 | 0.074 |
| Caste | *M. flaviceps* | Crop |  | wUnifrac | 0.23576 | 0.04196 * | 0.3237 | 0.037 | 2.7641 | 0.129 |
| Caste | *M. flaviceps* | Crop |  | Bray-Curtis | 0.08803 | 0.0959 | 0.1589 | 0.127 | 0.0389 | 0.849 |
| Caste | *M. flaviceps* | Midgut |  | wUnifrac | 0.24015 | 0.056 | 0.2421 | 0.075 | 4.2008 | 0.079 |
| Caste | *M. flaviceps* | Midgut |  | Bray-Curtis | 0.1717 | 0.122 | 0.09951 | 0.093 | 0.142 | 0.696 |
| Caste | *M. mimicus* | Crop |  | wUnifrac | 0.07423 | 0.2537 | 0.052 | 0.229 | 0.068 | 0.795 |
| Caste | *M. mimicus* | Crop |  | Bray-Curtis | 0.16573 | 0.01299 * | 0.2459 | 0.039 | 0.1643 | 0.709 |
| Caste | *M. mimicus* | Midgut |  | wUnifrac | 0.9171 | 0.04692 | -0.2407 | 0.97 | 0 | 0.97 |
| Caste | *M. mimicus* | Midgut |  | Bray-Curtis | 0.16654 | 0.4236 | -0.05556 | 0.684 | 7.00E-04 | 0.943 |
| Caste | *M. wheeleri* | Crop |  | wUnifrac | 0.07386 | 0.4186 | 0.05526 | 0.253 | 0.1934 | 0.663 |
| Caste | *M. wheeleri* | Crop |  | Bray-Curtis | 0.07816 | 0.7043 | 0.0125 | 0.478 | 0.7666 | 0.4 |
| Caste | *M. wheeleri* | Midgut |  | wUnifrac | 0.01884 | 0.933 | -0.07273 | 0.658 | 0.6524 | 0.471 |
| Caste | *M. wheeleri* | Midgut |  | Bray-Curtis | 0.07211 | 0.437 | 0.03818 | 0.323 | 0.1831 | 0.672 |
| Organ | All species |  | - | wUnifrac | 0.47863 | 0.000999 *** | - | - | 0.4027 | 0.538 |

| Organ | All species | - | Bray-Curtis | 0.20324 | 0.000999 *** | - | - | 0.5646 | 0.451 |
| --- | --- | --- | --- | --- | --- | --- | --- | --- | --- |
| Organ | *M. depilis* | - | wUnifrac | 0.06324 | 0.06394 | 0.08531 | 0.025 | 0.0774 | 0.78 |
| Organ | *M. depilis* | - | Bray-Curtis | 0.08565 | 0.002997 ** | 0.1169 | 0.014 | 3.5111 | 0.064 |
| Organ | *M. mexicanus* | - | wUnifrac | 0.22309 | 0.000999 *** | 0.3528 | 0.001 | 28.092 | 0.001 *** |
| Organ | *M. mexicanus* | - | Bray-Curtis | 0.21866 | 0.000999 *** | 0.4174 | 0.001 | 39.265 | 0.001 *** |
| Organ | *M. mimicus* | - | wUnifrac | 0.2297 | 0.002997 ** | 0.3308 | 0.005 | 0.6582 | 0.419 |
| Organ | *M. mimicus* | - | Bray-Curtis | 0.17693 | 0.000999 *** | 0.1105 | 0.129 | 18.422 | 0.001 *** |
| Organ | *M. wheeleri* | - | wUnifrac | 0.23282 | 0.001998 ** | 0.2151 | 0.011 | 3.3169 | 0.097 |
| Organ | *M. wheeleri* | - | Bray-Curtis | 0.06627 | 0.008991 ** | 0.1866 | 0.002 | 6.7312 | 0.022 * |
| Organ | *M. arenarius* | - | wUnifrac | 0.0219 | 0.3956 | -0.007101 | 0.37 | 0.0931 | 0.774 |
| Organ | *M. arenarius* | - | Bray-Curtis | 0.03102 | 0.3047 | -0.0003279 | 0.364 | 0.1082 | 0.739 |
| Organ | *M. flaviceps* | - | wUnifrac | 0.01278 | 0.8132 | -0.04012 | 0.909 | 0.2077 | 0.644 |
| Organ | *M. flaviceps* | - | Bray-Curtis | 0.03619 | 0.6713 | -0.01499 | 0.583 | 0.9539 | 0.348 |

Table S5. Summary of PERMANOVA, ANOSIM, and PERMDISP results using weighted UniFrac and Bray-Curtis distance matrices to assess the effect of collection time in replete crop and midgut samples of *M. mexicanus* and *M. depilis*.

| **Factor** | **Species** | **Organ** | **Distance Matrix** | **PERMANOVA R²** | **PERMANOVA p-value** | **ANOSIM R** |  | **ANOSIM p-value** | **PERMDISP F PERMDISP p-value** |
| --- | --- | --- | --- | --- | --- | --- | --- | --- | --- |
| Year | *M. depilis* | Crop | wUnifrac | 0.47145 | 0.000999 *** |  | 0.9178 | 0.002 | 2.1696 0.184 |
| Year | *M. depilis* | Crop | Bray-Curtis | 0.29525 | 0.000999 *** |  | 0.9729 | 0.001 | 0.3299 0.593 |
|  |  |  |  |  |  |  |  |  | PERMDISP was not evaluated because the July 2025 |
|  |  |  |  |  |  |  |  |  | group contained only one sample (n = 1), preventing |
| Year | *M. depilis* | Midgut | wUnifrac | 0.52155 | 0.0979 |  | 1 | 0.103 | estimation of within-group dispersion. |
|  |  |  |  |  |  |  |  |  | PERMDISP was not evaluated because the July 2025 |
|  |  |  |  |  |  |  |  |  | group contained only one sample (n = 1), preventing |
| Year | *M. depilis* | Midgut | Bray-Curtis | 0.38437 | 0.095 |  | 1 | 0.095 | estimation of within-group dispersion. |
| Year | *M. mexicanus* | Crop | wUnifrac | 0.08311 | 0.07493 |  | 0.4081 | 0.001 | 1.9661 0.121 |
| Year | *M. mexicanus* | Crop | Bray-Curtis | 0.31942 | 0.000999 *** |  | 0.6174 | 0.001 | 18.907 0.001 *** |
| Year | *M. mexicanus* | Midgut | wUnifrac | 0.53698 | 0.000999 *** |  | 0.7955 | 0.001 | 0.3115 0.749 |
| Year | *M. mexicanus* | Midgut | Bray-Curtis | 0.47642 | 0.000999 *** |  | 0.772 | 0.001 | 0.758 0.455 |
| Year 2023 vs 2025 M. depilis Crop wUnifrac 0.47145 0.000999 *** 0.9178 0.002 2.1696 0.184 | | | | | | | | | |
|  |  |  |  |  |  |  |  |  | PERMDISP was not evaluated because the July |
|  |  |  |  |  |  |  |  |  | 2025 group contained only one sample (n = 1), |
| Year 2023 vs 2025 | M. depilis | Midgut | wUnifrac | 0.52155 | 0.0979 |  | 1 | 0.103 | preventing estimation of within-group dispersion. |

Table S6. Shannon diversity comparisons between replete and worker castes within *Myrmecocystus* species using Wilcoxon rank sum tests. The raw p-values were adjusted for multiple comparisons using Benjamini Hochberg false discovery rate (FDR). FDR-adjusted p-values < 0.05 were considered statistically significant.

**Species	Raw p-value	FDR-adj p-value Significance**

***M. arenarius***	0.0132	0.0264	*

***M. depilis***	0.401	0.481	ns

***M. flaviceps***	0.00142	0.00453	**

***M. mexicanus***	0.00151	0.00453	**

***M. mimicus***	0.182	0.274	ns

***M. wheeleri***	0.724	0.724	ns

Table S7. Summary of *Myrmecocystus* species pairwise comparison using PERMANOVA, ANOSIM, and PERMDISP with weighted UniFrac and Bray-Curtis distance matrices.

| **Species 1** | **Species 2** | **Organ** | **Distance Matrix** | **PERMANOVA R²** | **PERMANOVA p-value** | **ANOSIM R** | **ANOSIM p-value** | **PERMDISP F** |  | **PERMDISP p-value** |
| --- | --- | --- | --- | --- | --- | --- | --- | --- | --- | --- |
| *M. mexicanus* | *M. depilis* | crop | wUnifrac | 0.16805 | 0.001 *** | 0.5194 | 0.001 |  | 21.926 | 0.001 *** |
| *M. mexicanus* | *M. depilis* | crop | Bray-Curtis | 0.15604 | 0.001 *** | 0.6183 | 0.001 |  | 22.655 | 0.001 *** |
| *M. mexicanus* | *M. wheeleri* | crop | wUnifrac | 0.66755 | 0.001 *** | 0.9925 | 0.001 |  | 3.5925 | 0.055 |
| *M. mexicanus* | *M. wheeleri* | crop | Bray-Curtis | 0.21263 | 0.001 *** | 0.9837 | 0.001 |  | 18.435 | 0.002 *** |
| *M. mexicanus* | *M. flaviceps* | crop | wUnifrac | 0.60741 | 0.001 *** | 0.9389 | 0.001 |  | 8.0189 | 0.005 ** |
| *M. mexicanus* | *M. flaviceps* | crop | Bray-Curtis | 0.22154 | 0.001 *** | 0.9788 | 0.001 |  | 18.947 | 0.001 *** |
| *M. depilis* | *M. wheeleri* | crop | wUnifrac | 0.57464 | 0.001 *** | 0.8593 | 0.001 |  | 4.1937 | 0.05* |
| *M. depilis* | *M. wheeleri* | crop | Bray-Curtis | 0.10559 | 0.001 *** | 0.5831 | 0.001 |  | 4.7423 | 0.036 * |
| *M. depilis* | *M. flaviceps* | crop | wUnifrac | 0.48896 | 0.001 *** | 0.7422 | 0.001 |  | 0.4383 | 0.497 |
| *M. depilis* | *M. flaviceps* | crop | Bray-Curtis | 0.10996 | 0.001 *** | 0.5732 | 0.001 |  | 3.0956 | 0.074 |
| *M. wheeleri* | *M. flaviceps* | crop | wUnifrac | 0.0211 | 0.639 | -0.0318 | 0.853 |  | 0.5516 | 0.457 |
| *M. wheeleri* | *M. flaviceps* | crop | Bray-Curtis | 0.03777 | 0.528 | 0.01115 | 0.368 |  | 0.354 | 0.578 |
| *M. mexicanus* | *M. arenarius* | crop | wUnifrac | 0.39394 | 0.001 *** | 0.7923 | 0.001 |  | 44.338 | 0.001 *** |
| *M. mexicanus* | *M. arenarius* | crop | Bray-Curtis | 0.26171 | 0.001 *** | 0.9705 | 0.001 |  | 17.223 | 0.001 *** |
| *M. mexicanus* | *M. mimicus* | crop | wUnifrac | 0.38509 | 0.001 *** | 0.7982 | 0.001 |  | 9.2815 | 0.007 ** |
| *M. mexicanus* | *M. mimicus* | crop | Bray-Curtis | 0.29634 | 0.001 *** | 0.9639 | 0.001 |  | 15.174 | 0.001 *** |
| *M. depilis* | *M. arenarius* | crop | wUnifrac | 0.22694 | 0.001 *** | 0.3677 | 0.001 |  | 5.0218 | 0.042 * |
| *M. depilis* | *M. arenarius* | crop | Bray-Curtis | 0.13905 | 0.001 *** | 0.582 | 0.001 |  | 0.0386 | 0.838 |
| *M. depilis* | *M. mimicus* | crop | wUnifrac | 0.23828 | 0.001 *** | 0.4241 | 0.001 |  | 1.2747 | 0.276 |
| *M. depilis* | *M. mimicus* | crop | Bray-Curtis | 0.14193 | 0.001 *** | 0.5004 | 0.001 |  | 2.3827 | 0.135 |
| *M. wheeleri* | *M. arenarius* | crop | wUnifrac | 0.15186 | 0.022 ** | 0.06576 | 0.09 |  | 10.243 | 0.009 ** |
| *M. wheeleri* | *M. arenarius* | crop | Bray-Curtis | 0.05902 | 0.026 * | 0.1081 | 0.043 |  | 2.2246 | 0.148 |
| *M. wheeleri* | *M. mimicus* | crop | wUnifrac | 0.66596 | 0.001 *** | 0.9386 | 0.001 |  | 0.4386 | 0.496 |
| *M. wheeleri* | *M. mimicus* | crop | Bray-Curtis | 0.13262 | 0.001 *** | 0.5964 | 0.001 |  | 9.1419 | 0.006 ** |
| *M. flaviceps* | *M. arenarius* | crop | wUnifrac | 0.11262 | 0.045 * | 0.06282 | 0.092 |  | 4.1589 | 0.048 * |
| *M. flaviceps* | *M. arenarius* | crop | Bray-Curtis | 0.08662 | 0.002 ** | 0.3273 | 0.002 |  | 1.6227 | 0.216 |
| *M. flaviceps* | *M. mimicus* | crop | wUnifrac | 0.58608 | 0.001 *** | 0.8592 | 0.001 |  | 0.0492 | 0.818 |
| *M. flaviceps* | *M. mimicus* | crop | Bray-Curtis | 0.15916 | 0.001 *** | 0.8274 | 0.001 |  | 7.5814 | 0.008 ** |
| *M. arenarius* | *M. mimicus* | crop | wUnifrac | 0.31535 | 0.001 *** | 0.4971 | 0.001 |  | 8.0179 | 0.011 * |
| *M. arenarius* | *M. mimicus* | crop | Bray-Curtis | 0.12389 | 0.001 *** | 0.308 | 0.001 |  | 0.9986 | 0.345 |

| *M. mexicanus* | *M. depilis* | midgut | wUnifrac | 0.09258 | 0.001 *** | 0.281 | 0.001 | 3.7849 | 0.061 |
| --- | --- | --- | --- | --- | --- | --- | --- | --- | --- |
| *M. mexicanus* | *M. depilis* | midgut | Bray-Curtis | 0.05563 | 0.001 *** | 0.2503 | 0.001 | 0.0264 | 0.878 |
| *M. mexicanus* | *M. wheeleri* | midgut | wUnifrac | 0.38086 | 0.001 *** | 0.8786 | 0.001 | 8.9623 | 0.005** |
| *M. mexicanus* | *M. wheeleri* | midgut | Bray-Curtis | 0.10446 | 0.001 *** | 0.6447 | 0.001 | 3.2511 | 0.082 |
| *M. mexicanus* | *M. flaviceps* | midgut | wUnifrac | 0.53571 | 0.001 *** | 0.9414 | 0.001 | 0.0101 | 0.927 |
| *M. mexicanus* | *M. flaviceps* | midgut | Bray-Curtis | 0.11003 | 0.001 *** | 0.9112 | 0.001 | 13.232 | 0.001 *** |
| *M. depilis* | *M. wheeleri* | midgut | wUnifrac | 0.28445 | 0.001 *** | 0.4731 | 0.001 | 0.0022 | 0.962 |
| *M. depilis* | *M. wheeleri* | midgut | Bray-Curtis | 0.19844 | 0.001 *** | 0.5775 | 0.001 | 0.9354 | 0.353 |
| *M. depilis* | *M. flaviceps* | midgut | wUnifrac | 0.44815 | 0.001 *** | 0.5701 | 0.001 | 0.5315 | 0.459 |
| *M. depilis* | *M. flaviceps* | midgut | Bray-Curtis | 0.16959 | 0.001 *** | 0.5482 | 0.001 | 3.5115 | 0.074 |
| *M. wheeleri* | *M. flaviceps* | midgut | wUnifrac | 0.22548 | 0.005 ** | 0.2481 | 0.004 | 1.3344 | 0.249 |
| *M. wheeleri* | *M. flaviceps* | midgut | Bray-Curtis | 0.11349 | 0.001 *** | 0.5241 | 0.001 | 3.4899 | 0.074 |
| *M. mexicanus* | *M. arenarius* | midgut | wUnifrac | 0.21565 | 0.001 *** | 0.6516 | 0.001 | 69.511 | 0.001*** |
| *M. mexicanus* | *M. arenarius* | midgut | Bray-Curtis | 0.11946 | 0.001 *** | 0.7258 | 0.001 | 5.7272 | 0.019 * |
| *M. mexicanus* | *M. mimicus* | midgut | wUnifrac | 0.1457 | 0.001 *** | 0.5837 | 0.001 | 1.1519 | 0.282 |
| *M. mexicanus* | *M. mimicus* | midgut | Bray-Curtis | 0.08658 | 0.001 *** | 0.4287 | 0.001 | 12.579 | 0.001 *** |
| *M. depilis* | *M. arenarius* | midgut | wUnifrac | 0.11655 | 0.02 * | 0.2147 | 0.003 | 3.3059 | 0.078 |
| *M. depilis* | *M. arenarius* | midgut | Bray-Curtis | 0.18424 | 0.001 *** | 0.5764 | 0.001 | 1.5576 | 0.21 |
| *M. depilis* | *M. mimicus* | midgut | wUnifrac | 0.19254 | 0.007 ** | 0.3532 | 0.009 | 0.8739 | 0.374 |
| *M. depilis* | *M. mimicus* | midgut | Bray-Curtis | 0.28518 | 0.001 *** | 0.5247 | 0.001 | 2.6798 | 0.113 |
| *M. wheeleri* | *M. arenarius* | midgut | wUnifrac | 0.0614 | 0.135 | 0.07551 | 0.092 | 7.5664 | 0.017* |
| *M. wheeleri* | *M. arenarius* | midgut | Bray-Curtis | 0.05093 | 0.03 * | -0.0121 | 0.556 | 0.136 | 0.721 |
| *M. wheeleri* | *M. mimicus* | midgut | wUnifrac | 0.26815 | 0.005** | 0.2313 | 0.021 | 3.7957 | 0.066 |
| *M. wheeleri* | *M. mimicus* | midgut | Bray-Curtis | 0.11373 | 0.004 ** | -0.09433 | 0.799 | 15.657 | 0.001*** |
| *M. flaviceps* | *M. arenarius* | midgut | wUnifrac | 0.24887 | 0.003 *** | 0.2043 | 0.013 | 10.748 | 0.006** |
| *M. flaviceps* | *M. arenarius* | midgut | Bray-Curtis | 0.11877 | 0.001*** | 0.5516 | 0.001 | 1.2037 | 0.282 |
| *M. flaviceps* | *M. mimicus* | midgut | wUnifrac | 0.60958 | 0.001*** | 0.8272 | 0.001 | 0.2234 | 0.655 |
| *M. flaviceps* | *M. mimicus* | midgut | Bray-Curtis | 0.25226 | 0.001*** | 0.7326 | 0.001 | 45.929 | 0.001*** |
| *M. arenarius* | *M. mimicus* | midgut | wUnifrac | 0.10964 | 0.069 | 0.04652 | 0.2 | 13.932 | 0.003** |
| *M. arenarius* | *M. mimicus* | midgut | Bray-Curtis | 0.11244 | 0.003 ** | -0.164 | 0.962 | 14.403 | 0.001*** |

Table 8a. GLMM sample size on decontam non-rarefied data

|  | **Replete** | **Worker** | **Crop** | **Midgut** |
| --- | --- | --- | --- | --- |
| ***M. arenarius*** | 37 | 29 | 30 | 36 |
| ***M. depilis*** | 37 | 27 | 32 | 31 |
| ***M. flaviceps*** | 17 | 11 | 15 | 13 |
| ***M. mexicanus*** | 104 | 68 | 88 | 84 |
| ***M. mimcus*** | 27 | 20 | 23 | 24 |
| ***M. wheeleri*** | 18 | 10 | 14 | 14 |

Table 8b. GLMM colony sample size on decontaminated, non-rarefied data.

| **Colony** | **Repletes** | **Workers** |
| --- | --- | --- |
| A1 | 10 | 12 |
| A2 | 6 | 4 |
| A3 | 9 | 4 |
| A4 | 7 | 5 |
| A5 | 5 | 4 |
| D1 | 19 | 9 |
| D2 | 8 | 9 |
| D3 | 10 | 9 |
| M1 | 17 | 10 |
| M2 | 10 | 10 |
| V1 | 16 | 7 |
| V2 | 1 | 4 |
| W1 | 10 | 0 |
| W2 | 8 | 10 |
| X1 | 15 | 7 |
| X2 | 12 | 9 |
| X3 | 9 | 6 |
| X4 | 10 | 6 |
| X7 | 38 | 10 |
| X8 | 8 | 10 |
| X9 | 2 | 9 |
| X10 | 10 | 9 |

Table 9. Type II Wald chi-square tests from a Gaussian generalized linear mixed-effects model (GLMM) examining the effects of host species, digestive organ, and caste on microbiome Shannon diversity in Myrmecocystus ants.

| **Predictor** | **Chisq** | **df** | **Pr (>Chisq)** |
| --- | --- | --- | --- |
| Species | 29.8937 | 5 | 1.548e-05 *** |
| Organ | 113.4394 | 1 | < 2.2e-16 *** |
| Caste | 2.7996 | 1 | 0.09429 . |
| Species:Organ | 101.0615 | 5 | < 2.2e-16 *** |

Table 10. Pairwise comparisons of Shannon diversity between crop and midgut microbiomes within *Myrmecocystus* species, based on estimated marginal means from a Gaussian generalized linear mixed-effects model (GLMM). Estimates represent the difference in Shannon diversity between the crop and midgut (crop − midgut), with negative values indicating greater diversity in the midgut.

Comparisons were averaged over caste.

| **Species** | **Estimate (crop - SE** |  | **z-ratio** |  | **p-value** |
| --- | --- | --- | --- | --- | --- |
| *M. arenarius* | -0.0283 | 0.184 | -0.154 | | 0.8779 |
| *M. depilis* | -0.0352 | 0.188 | -0.188 | | 0.8512 |
| *M. flaviceps* | -0.1033 | 0.286 | -0.361 | | 0.7183 |
| *M. mexicanus* | -1.6298 | 0.113 | -14.378 | | <0.0001 |
| *M. mimicus* | -0.5938 | 0.215 | -2.756 | | 0.0058 |
| *M. wheeleri* | -0.0875 | 0.279 | -0.314 | | 0.7538 |

Table 11. Pairwise comparisions of Shannon diversity across six *Myrmecocystus* species within the crop or midgut based on estimated marginal means from a GLMM. Estimates represent differences in Shannon diversity between species (Species 1 − Species 2). Estimates, standard errors (SE), z-ratios, and Tukey-adjusted p-values are reported. Comparisons were averaged over caste.

| **Species 1** | **Species 2** | **Organ** | **Estimate** | **SE** |  | **z-ratio** |  | **p-value** |
| --- | --- | --- | --- | --- | --- | --- | --- | --- |
| *M. arenarius* | *M. depilis* | crop | -0.34815 | 0.302 | | -1.153 | | 0.8591 |
| *M. arenarius* | *M. flaviceps* | crop | -0.98743 | 0.363 | | -2.717 | | 0.0719 |
| *M. arenarius* | *M. mexicanus* | crop | 1.01778 | 0.244 | | 4.173 | | 0.0004 |
| *M. arenarius* | *M. mimicus* | crop | -0.00417 | 0.338 | | -0.012 | | 1 |
| *M. arenarius* | *M. wheeleri* | crop | -0.84234 | 0.362 | | -2.325 | | 0.1838 |
| *M. depilis* | *M. flaviceps* | crop | -0.63928 | 0.379 | | -1.689 | | 0.5392 |
| *M. depilis* | *M. mexicanus* | crop | 1.36592 | 0.266 | | 5.138 | | <0.0001 |
| *M. depilis* | *M. mimicus* | crop | 0.34398 | 0.355 | | 0.97 | | 0.9275 |
| *M. depilis* | *M. wheeleri* | crop | -0.49419 | 0.378 | | -1.309 | | 0.7804 |
| *M. flaviceps* | *M. mexicanus* | crop | 2.00521 | 0.333 | | 6.015 | | <0.0001 |
| *M. flaviceps* | *M. mimicus* | crop | 0.98326 | 0.408 | | 2.411 | | 0.1523 |
| *M. flaviceps* | *M. wheeleri* | crop | 0.14509 | 0.428 | | 0.339 | | 0.9994 |
| *M. mexicanus* | *M. mimicus* | crop | -1.02194 | 0.306 | | -3.338 | | 0.0109 |
| *M. mexicanus* | *M. wheeleri* | crop | -1.86012 | 0.333 | | -5.59 | | <0.0001 |
| *M. mimicus* | *M. wheeleri* | crop | -0.83817 | 0.407 | | -2.058 | | 0.3094 |
| *M. arenarius* | *M. depilis* | midgut | -0.35512 | 0.296 | | -1.199 | | 0.8377 |
| *M. arenarius* | *M. flaviceps* | midgut | -1.06244 | 0.379 | | -2.8 | | 0.0575 |
| *M. arenarius* | *M. mexicanus* | midgut | -0.58379 | 0.236 | | -2.47 | | 0.133 |
| *M. arenarius* | *M. mimicus* | midgut | -0.56975 | 0.331 | | -1.721 | | 0.5176 |
| *M. arenarius* | *M. wheeleri* | midgut | -0.90154 | 0.357 | | -2.525 | | 0.1168 |
| *M. depilis* | *M. flaviceps* | midgut | -0.70732 | 0.4 | | -1.77 | | 0.4852 |
| *M. depilis* | *M. mexicanus* | midgut | -0.22867 | 0.268 | | -0.855 | | 0.957 |
| *M. depilis* | *M. mimicus* | midgut | -0.21462 | 0.354 | | -0.606 | | 0.9906 |
| *M. depilis* | *M. wheeleri* | midgut | -0.54642 | 0.379 | | -1.443 | | 0.7005 |
| *M. flaviceps* | *M. mexicanus* | midgut | 0.47865 | 0.357 | | 1.341 | | 0.7622 |

| *M. flaviceps* | *M. mimicus* | midgut | 0.49269 | 0.426 | 1.157 | 0.8572 |
| --- | --- | --- | --- | --- | --- | --- |
| *M. flaviceps* | *M. wheeleri* | midgut | 0.1609 | 0.447 | 0.36 | 0.9992 |
| *M. mexicanus* | *M. mimicus* | midgut | 0.01405 | 0.305 | 0.046 | 1 |
| *M. mexicanus* | *M. wheeleri* | midgut | -0.31775 | 0.333 | -0.953 | 0.9324 |
| *M. mimicus* | *M. wheeleri* | midgut | -0.33179 | 0.406 | -0.817 | 0.9645 |
